# Molecule-resolved inference reveals the spatial organization of transcriptional programs

**DOI:** 10.64898/2026.10.29.762960

**Authors:** Yichen Si, ChangHee Lee, Weiqiu Cheng, Michael Sweeney, Jeong H. Yun, Myungjin Kim, Goo Jun, Jun Hee Lee, Hyun Min Kang

**Author notes:** Correspondence should be addressed to H.M.K., J.H.L., and Y.S.

## Abstract

Transcriptional programs encode cellular identity and physiological activity, linking gene regulation to cellular phenotypes. Although spatial transcriptomics measures individual RNA molecules across intact tissues, most analyses aggregate them into cells or pixels, obscuring molecular level functional dynamics. We introduce FICTURE2, a statistical model that infers the transcriptional program from which each individual molecule is generated, using gene identity and spatial context alone without segmentation or binning. FICTURE2 learns programs with distinct spatial distributions, ranging from millimeter-scale continuous transition to micron-scale subcellular localization, the coexisting patterns of which then reveal both cellular identity and physiological activity with unified interpretations. FICTURE2 supports both unsupervised and reference-guided inference and is orders of magnitude more efficient than alternative approaches, enabling systematic analysis of large-scale spatial datasets as we demonstrate in data across diverse tissue types profiled by various spatial technologies.

## Introduction

Spatial transcriptomics can now locate millions to billions of individual RNA transcripts across intact tissues at subcellular resolution, revealing molecular organization inaccessible to bulk or dissociated single-cell transcriptomics.^1^ By preserving the native spatial context of gene expression, recent sequencing- and imaging-based technologies enable direct investigation of how transcripts are localized within cells, how physiological activities are distributed across tissues, and how these patterns change during development and disease.^2–10^ Yet the resolution of biological analysis has lagged behind that of measurement. Most analytical approaches continue to aggregate transcripts into spatial bins, segmented cells, or regions of interest. Cell segmentation remains error-prone in densely packed tissues and across complex cell morphologies.^11–13^ More fundamentally, even perfect segmentation collapses the spatial arrangement of a cell’s transcripts into a single expression profile, erasing subcellular organization within individual cells.

Aggregating transcripts at the cellular level also obscures how multiple biological processes coexist and interact across spatial scales. Although cells are fundamental biological units, their transcriptomes do not necessarily represent single, discrete functional states.^14^ Individual cells simultaneously express multiple transcriptional programs, each a coherent pattern of gene expression reflecting a cellular identity or activity, such as differentiation, proliferation, metabolism, and responses to environmental stimuli.^15^ A proliferating cell retains its lineage identity while expressing cell-cycle programs, just as a differentiated cell responding to injury may preserve its differentiation program while acquiring additional stress- or injury-associated programs. Single-cell transcriptomic factorization has demonstrated that identity and activity programs can coexist within individual cells.^16^ However, they cannot resolve how co-occurring programs are localized within individual cells or coordinated across neighboring cells and tissue microenvironments.

These considerations suggest that individual RNA molecules, rather than predefined cells or spatial bins, should serve as the primary units of spatial inference, with transcriptional programs serving as the units of biological interpretation. Based on this insight, we introduce FICTURE2, which infers the transcriptional programs from which each molecule is originated, leveraging the two fundamental observations per molecule: its gene identity and its location within the tissue. Gene identity constrains the programs to which a transcript may belong, whereas its spatial neighborhood indicates which programs are active nearby. FICTURE2 resolves co-occurring programs even when their transcripts are intermingled within the same cell or even the same pixel. These molecular assignments can subsequently be aggregated into cells or tissue regions, allowing biological organization to be examined across multiple spatial scales without requiring cell boundaries as a prerequisite for inference.

FICTURE2 establishes individual RNA molecules as the units of spatial inference to resolve transcriptional programs across intact tissues. By integrating each transcript’s gene identity with the local distribution of active programs, FICTURE2 probabilistically assigns individual molecules to programs without assuming that neighboring transcripts are generated from the same program. Program profiles can be learned from spatial transcriptomic data or from single-cell references, allowing the dual interpretation of cellular identities and overlapping transcriptional programs within a common framework. Across diverse tissues, we demonstrate that molecular-level inference reveals the persistence of differentiation programs alongside injury-response programs during wound repair, emerging fiber-type and neuromuscular-junction programs within developing skeletal muscle, and subcellularly localized gene expression programs within colorectal tumor cells. We further identify activity-associated programs shared across distinct cell lineages and tumors and detect disease-associated expression changes in fibrotic lung that are not explained by changes in cellular composition alone. Supporting imaging- and sequencing-based spatial transcriptomics with near-linear computational scaling to datasets across billions of transcripts, FICTURE2 provides a framework for investigating tissue organization through the spatial distribution and coexistence of transcriptional programs rather than discrete cellular identities alone.

## Results

### FICTURE2 resolves transcriptional programs at molecular resolution

Our original motivation for developing FICTURE was to overcome the limitations of cell segmentation by inferring cellular identities directly from spatial transcriptomic data.^17^ After applying it to a wider range of tissues in increasingly large data collections,^18^ however, we observed that unsupervised factorizations learn interpretable and reproducible transcriptional programs that do not align with known discrete cell-types or conform to cellular boundaries. Revising previous statistical interpretation that the overlapping and sometimes diffused spatial distributions reflect imperfect cell-type inference, here we reconsider the fundamental unit of spatial inference and developed FICTURE2 to infer the source program for each individual molecule. The key biological insight encoded in FICTURE2 that distinguishes it from existing methods is that a cell may simultaneously express multiple transcriptional programs, therefore neighboring transcripts, even those located within half a micron, need not share the same program assignment. FICTURE2 realizes this principle by probabilistically assigning each molecule to transcriptional programs present in the neighborhood based on its gene identity and the programs’ spatial distribution (Fig. 1A). FICTURE assigns a common factor to transcripts within each microscopic pixel; FICTURE2 further resolves programs jointly activated within a single cell. Combined with the flexibility of learning the programs from ST data or from reference single cell data, our extended framework offers dual interpretations of the molecular observations in terms of both conventional cell types and combinations of transcriptional programs.

**Figure 1.**
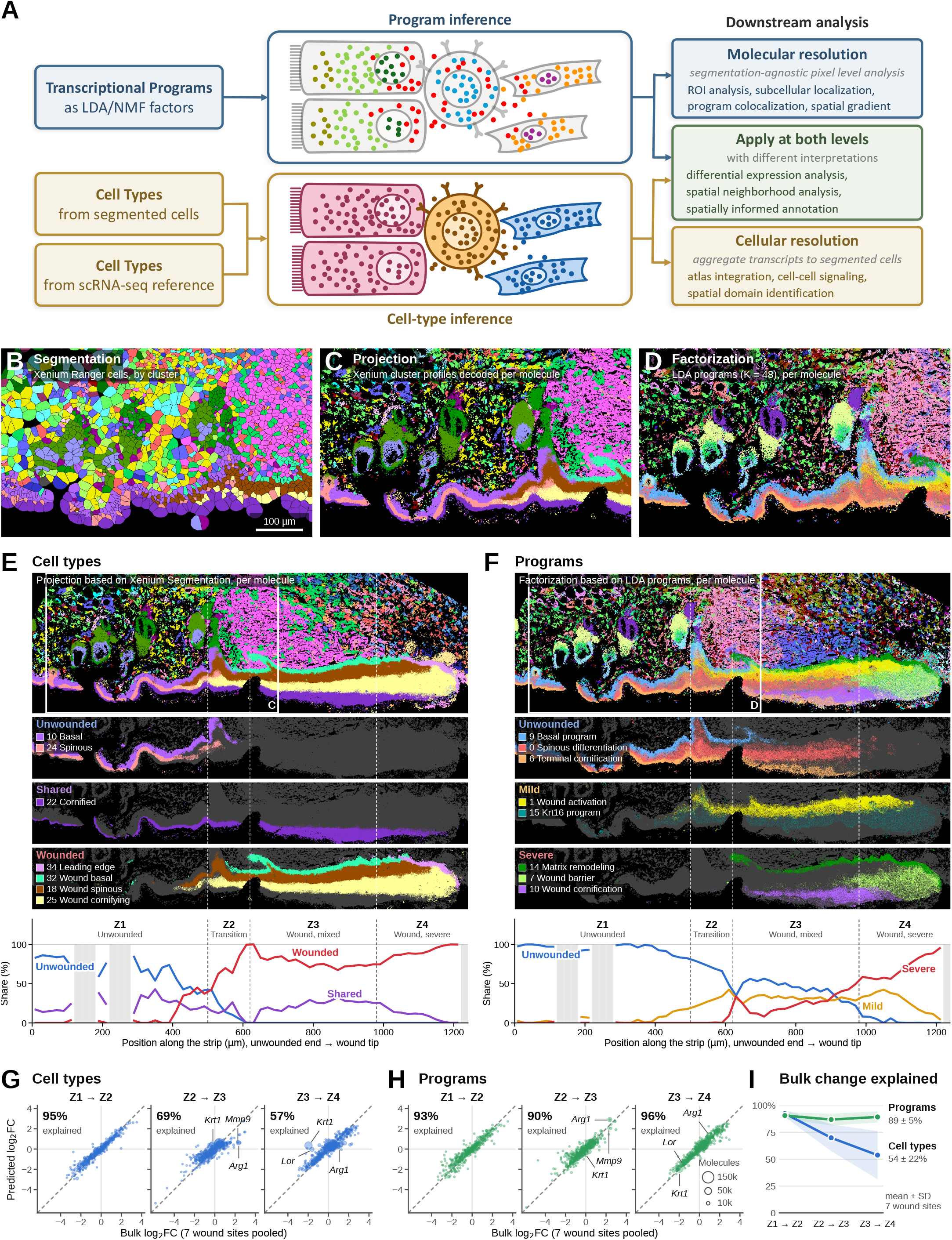
Molecular-resolution program inference on gradual wound responses (A) Schematic view of dual-level inference of FICTURE2. FICTURE2 takes a pseudobulk matrix (genes by factors) as an input to assign spatially resolved transcripts (or molecules) into cell types or transcriptional programs individually. In the cell-type inference mode (i.e. FICTURE1 mode), each column (factor) in the pseudobulk matrix is assumed to represent cell types, and transcripts within a specified distance (e.g. 0.5µm) are collapsed to share the same pixel level posterior inference. In the program inference mode (i.e. FICTURE2 mode), each column in the pseudobulk matrix is assumed to represent a transcriptional program, and the posterior distribution of programs are inferred for each transcript separately (see Methods for details), allowing transcripts located at the same position to be assigned to different transcriptional programs. **(B)** Segmented cells of mouse skin six days after wounding, colored by cell-type cluster generated by Xenium Ranger; the region is boxed in E and F. **(C)** The cluster profiles of B projected onto individual transcripts by FICTURE2, same region and colors. **(D)** 48 transcriptional programs learned from the same data by unsupervised LDA and assigned to individual transcripts by FICTURE2, same region. **(E)** Xenium Ranger cell types along an epidermal strip of wound site from unwounded skin to the wound tip (box, region of B–D), with unwounded, shared and wounded cell types highlighted (middle) and their shares among these seven cell types along the strip (bottom). Zones Z1–Z4 are defined from the program shares in F. **(F)** As in E, for transcriptional programs. Z2, Z3 and Z4 begin where the mild wound programs reach 20% and the severe wound programs reach **1**0% and 50%, respectively. **(G)** Observed bulk log2 fold change of each gene between adjacent zones versus the change predicted from cell-type composition with cell-type expression profiles held fixed; percentages, share of the bulk change explained. **(H)** As in G, predicted from program composition. **(I)** Share of the bulk change explained by programs and cell types, computed separately at each of the seven wound sites.

### Molecular inference reveals gradual wound responses obscured by cell-type classification

We examined a Xenium dataset of mouse skin collected six days after wounding (476 genes; 29.1 million transcripts) to compare cell-based annotation with molecular-level inference.^19^ Across the stratified epidermis, segmented cells have unrealistic shapes and cell-type assignments from Xenium Ranger clustering are fragmented (Fig. 1B). In contrast, FICTURE2 projection from the same cell-type profiles resolved continuous basal, spinous, and cornified epidermal layers, together with well-defined hair follicles and dermal structures (Fig. 1C). Thus, molecular-level inference markedly improved spatial annotation without changing the underlying cell-type profiles. We then learned 48 transcriptional programs directly from spatial transcriptomic data using latent Dirichlet allocation (LDA). Molecular-level assignment of these programs preserved tissue architecture while revealing physiological activities with spatial patterns beyond cell-type distribution (Fig. 1D).

Across a 1,240-µm epidermal strip divided into four zones extending from unwounded skin (Z1) to the wound tip (Z4), cell-type annotation showed an abrupt transition from normal to wound-associated keratinocytes (Fig. 1E). Unwounded basal and spinous cell types declined from accounting for 42% of keratinocyte transcripts at 500-520 µm to less than 0.5% beyond 600 µm, where wound-associated types dominated (72-89% across Z2-Z4). In the program view, however, basal, spinous differentiation, and terminal cornification programs persisted into the wounded epidermis, still accounting for 42% of transcripts in Z3 (declining from 56% to 22% across the zone), where unwounded cell-type labels had disappeared (Fig. 1F). Meanwhile, mild wound-response programs (*Aqp3*, *Areg*, *Sdc1*, *Krt16*) increased from approximately 5% in Z1 to 30% in Z2, whereas severe programs associated with matrix remodeling, barrier formation, and wound cornification increased from 2% in Z2 to 24% in Z3 and 67% in Z4 (Fig. 1F). These same contrasts between cell types and programs recurred across seven wound sites (Figs. S1A-S1C). Changes in program composition also aligned with the observed expression gradient of *Krt1* and *Arg1* within the wound-cornifying cell type (Figs. S1D and S1E).

Discrete cell types are insufficient for capturing the gene expression changes in the wound sites. For each of the seven wound sites and each zone, we predict the zone-level expression profile based on the cell-type and program compositions separately then ask if the changes in the compositions between zones explain the observed changes in gene expressions. Both approaches explained most expression changes from Z1 to Z2 (95% for cell types and 93% for programs; Figs. 1G and 1H). However, cell-type composition explained only 69% and 57% of changes from Z2 to Z3 and Z3 to Z4, respectively, compared with 90% and 96% for programs (Figs. 1G and 1H). This difference was consistent across individual wound sites, where programs explained 89 ± 5% of changes from Z3 to Z4, compared with 54 ± 22% for cell types (mean ± SD; Fig. 1I). Thus, wound progression involves the gradual remodeling of transcriptional programs while differentiation programs persist, a process largely obscured by discrete cell-type classification.

### Molecular inference resolves developmental and functional heterogeneity within skeletal myofibers

We next examined a whole P1 mouse pup profiled by Xenium Prime (5,010 genes), focusing on programs emerging during early development. Developing skeletal muscles consisting of multinucleated myofibers are challenging for cell segmentation. In neonatal mouse facial muscle (a region with 10.3 million transcripts), Xenium Ranger segmentation divided continuous myofibers into small, nucleus-centered fragments (Fig. 2A). Cell-type clustering identified the overall muscle architecture (Fig. 2B) but assigned 77.7% of labeled muscle transcripts to a single skeletal-myofiber cluster, blind to their developmental and functional heterogeneity (Fig. 2C and 2D).

**Figure 2.**
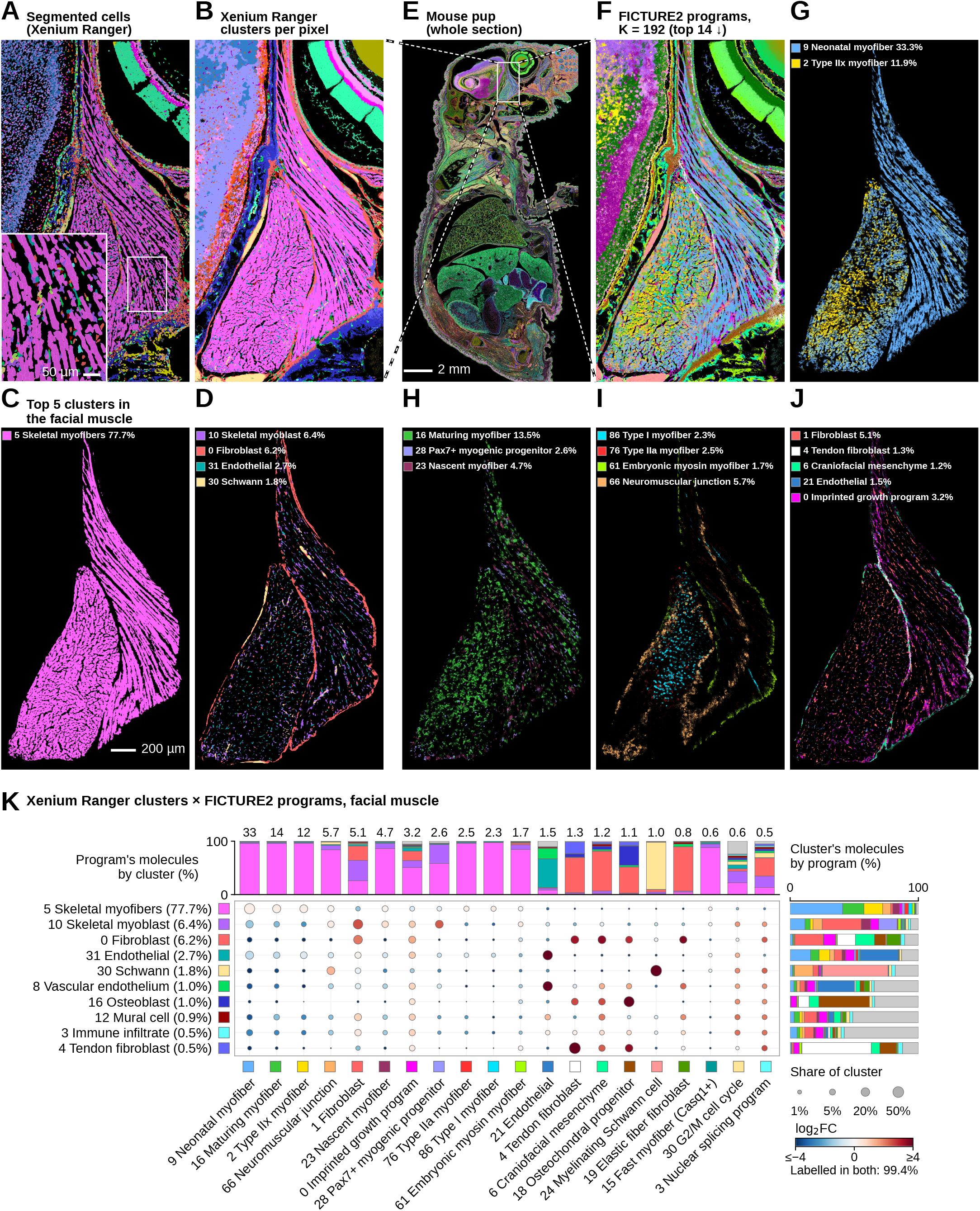
Molecule-level program analysis of neonatal mouse facial muscle. **(A)** Xenium Ranger segmented cells colored by cluster in the facial muscle of a neonatal mouse pup (field boxed in E); inset, myofiber segments are small fragments rather than whole fibers. **(B)** The same clusters assigned to individual transcripts by FICTURE2 from their expression profiles, without cell boundaries. **(C)** The largest cluster of B within the facial-muscle region; percentages are shares of the region’s labelled transcripts. The scale bar applies to A–D and F–J. **(D)** The next four abundant clusters. **(E)** Whole sagittal section colored by FICTURE2 program; the box marks the field of A, B and F. **(F)** FICTURE2 programs inferred from the whole section without segmentation; the 14 programs with the largest shares of the facial muscle are shown in G–J. **(G)** Neonatal and type IIx myofiber programs within the facial-muscle region, as in C. **(H)** Programs of developing myofibers and myogenic progenitors, as in G. **(I)** Fiber-type and localized myofiber programs, as in G; the neuromuscular-junction program forms discrete foci within the fiber bundles. **(J)** Stromal, vascular and imprinted growth programs, as in G. **(K)** Cross-tabulation of clusters (as in B) and programs (as in F) over the facial-muscle transcripts, for labels holding at least 0.5% of the region. Color, log2 ratio of observed to expected joint transcript mass; numbers above the bars, each program’s share of the region (%).

We inferred 192 transcriptional programs from the whole-section transcriptome and assigned individual molecules to these programs (Fig. 2E and 2F). Although the neonatal myofiber program (*Myh8*) remained dominant (33.3% of muscle transcripts), we detected distinct fiber-type programs, including type IIx (11.9%; *Myh1*), IIa (2.5%; *Myh2*), and I (2.3%; *Myh7*), indicating emerging fiber-type specification in immature muscle (Figs. 2G-2I, and S2A). Additional programs captured myofiber maturation (13.5%), nascent myofibers expressing *Myog* and *Mymk* (4.7%), Pax7-positive myogenic progenitors (2.6%), and embryonic myosin expression (1.7%) (Figs. 2H, 2I, and S2A). Imprinted growth-associated (3.2%) and surrounding stromal and vascular programs were also spatially resolved (Fig. 2J). Notably, a neuromuscular-junction (NMJ) program (5.7%), marked by *Chrna1*, *Chrnd*, *Colq*, and *Ache*, formed discrete foci within myofiber bundles, resolving specialized subcellular domains that were not distinguished by cell-type annotation (Figs. 2I and S2A).

These programs exhibited distinct gene signatures (Fig. S2A) and spatial distributions but intermingled within regions annotated as skeletal myofibers (Fig. 2F-I). Indeed, the single skeletal-myofiber cluster was distributed among neonatal (41.0%), maturing (16.6%), type IIx (14.7%), NMJ (6.1%), and nascent myofiber (5.2%) programs, among others (Figs. 2K and S2B). Thus, molecular-level inference distinguished emerging fiber-type identities and localized functional specialization within developing muscle that conventional cell-type annotation could not resolve.

### Molecular inference reveals subcellular organization and functional heterogeneity in colorectal cancer

In wound response and functional specialization, multiple programs activated in the same cell coexist in space, but sometimes transcripts from a program also show distinct subcellular localizations. We applied the same unsupervised FICTURE2 workflow to a published human colorectal cancer tissue section^20^ profiled by Xenium Prime 5K (5,001 genes; 281.8 million transcripts), identifying 48 transcriptional programs (Fig. 3A). These programs distinguished normal colonic epithelium (Fig. 3B) from malignant tissue and revealed substantial intratumoral heterogeneity lost in cell-type annotation (Fig. S3A). Four tumor-associated programs were distributed across malignant epithelial regions but were particularly prominent in the tumor core (Fig. 3C), collectively increasing from 17.6% of epithelial transcripts near the luminal surface to 52.2% in deeper regions (Fig. S3B). In contrast, neuroendocrine-like and inflammatory programs were enriched toward tumor surfaces (Fig. 3D), whereas a lipogenic program is enriched between the surface and the core (Fig. 3E). These programs exhibited distinct gene expression signatures, indicating substantial functional heterogeneity within the malignant epithelium (Fig. S3C).

**Figure 3.**
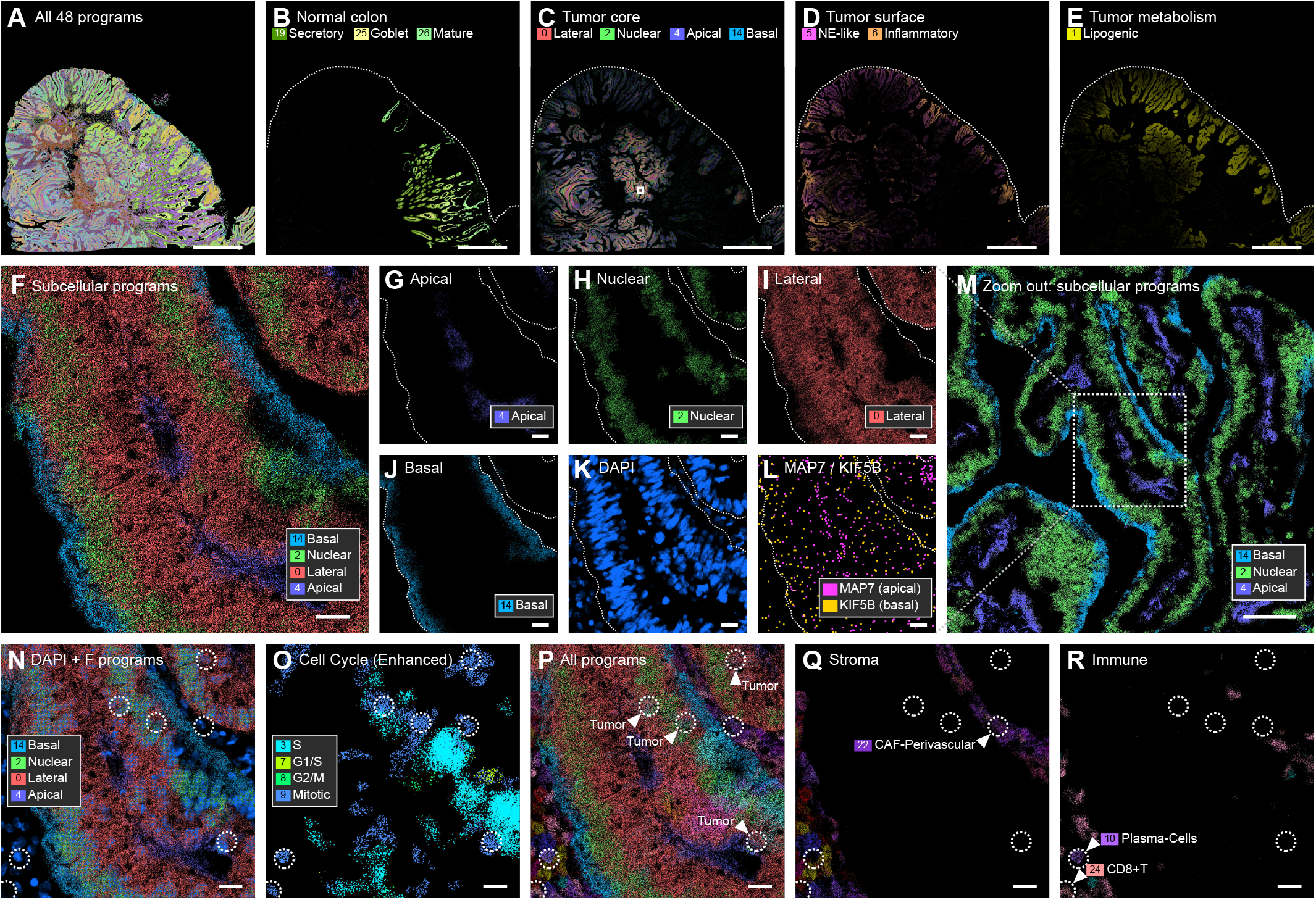
Molecular inference reveals subcellular organization and functional heterogeneity in colorectal cancer. **(A)** All 48 FICTURE2 programs in a human colorectal cancer section (Xenium Prime 5K). **(B)** Programs of normal colonic epithelium, confined to the normal crypts. Dotted line, luminal surface (also in C–E). **(C)** Tumor-core programs, present across the malignant epithelium and most prominent in the deep tumor (Fig. S3B). Box, region shown in F–M. **(D)** Neuroendocrine-like and inflammatory programs, enriched toward the tumor surface. **(E)** The lipogenic program, broadly distributed but relatively enriched near the surface. **(F)** Tumor glands (box in M) with each transcript colored by its subcellular program: the basal program lies next to the stroma and the apical program toward the gland lumen. **(G)** The apical program alone, concentrated toward the gland lumen; G–L show the same field. **(H)** The nuclear program alone, coinciding with the nuclear layer in K. **(I)** The lateral program alone. **(J)** The basal program alone, a thin layer along the gland border next to the stroma. **(K)** DAPI nuclear stain. **(L)** Individual transcripts of *MAP7* and *KIF5B*, markers of the apical and basal programs (Fig. S3C). **(M)** Wider view around F (box): the same basal-to-apical layering recurs across glands. **(N)** DAPI with the subcellular programs in the same field as F-L containing tumor epithelium, stroma and immune cells; N–R show the same field, with dotted circles highlighting mitotic cells. **(O)** Cell-cycle programs. **(P)** All programs; arrowheads, cell-cycle foci in tumor epithelium. **(Q)** Stromal programs; arrowhead, a cell-cycle focus within the perivascular fibroblast program. **(R)** Immune programs; arrowheads, cell-cycle foci within the plasma-cell and CD8+ T-cell programs (quantified in Fig. S3H).

Unexpectedly, the four prominent tumor-associated programs did not represent distinct malignant cell populations but instead exhibited striking subcellular organization within the same tumor glands (Fig. 3F). They occupied distinct apical (Fig. 3G), nuclear (Fig. 3H), lateral (Fig. 3I), and basal (Fig. 3J) regions, with the apical program concentrated toward glandular lumens and the basal program adjacent to surrounding stroma. Their distributions were supported by nuclear staining (Fig. 3K) and the differential localization of representative transcripts, such as *MAP7* and *KIF5B* (Fig. 3L). Similar polarized organization was observed across multiple tumor glands (Fig. 3M). Quantitative analysis confirmed the spatial separation of these programs along the stroma-to-lumen axis and nuclear-cytoplasm organization (Figs. S3D-S3F). Independent MAP7 immunohistochemistry images from antibody validation resources and the Human Protein Atlas further supported its preferential localization to the apical, luminal side of colorectal tumor glands (Fig. S3G).

Beyond subcellular polarity, FICTURE2 identified distinct cell-cycle programs corresponding to S, G1/S, G2/M, and mitotic states (Fig. 3N-3P), coexisting spatially with polarized epithelial programs (Fig. 3N). Notably, these proliferation-associated programs (Fig. 3O) were not restricted to malignant epithelial cells (Fig. 3P; mitotic phase cells highlighted in circles). The same programs were also detected in stromal (Fig. 3Q) and immune populations (Fig. 3R), superimposed on their distinct lineage-associated programs. Quantitative analysis confirmed their distribution across epithelial, stromal, lymphoid, and myeloid populations (Fig. S3H).

Together, these findings revealed multiple levels of molecular organization within the same tumor. Individual malignant epithelial cells contained distinct combinations of polarity-associated, cell-cycle, lipogenic, neuroendocrine-like, and inflammatory programs (Figs. S3I and S3J), with recurrent patterns of program coexistence across the malignant cell population (Fig. S3K). Furthermore, different tumor regions were surrounded by distinct stromal and immune programs (Fig. S3L), and their spatial abundances exhibited significant associations with specific tumor programs, even after accounting for tissue location (Fig. S3M). Thus, molecular inference revealed how multiple biological programs coexist within malignant cells and how their spatial distributions are associated with the surrounding tumor microenvironment.

### Joint molecular inference reveals conserved transcriptional programs across distinct tumors

We next leveraged FICTURE2’s computational advance and multi-sample capacity to identify transcriptional programs shared across diverse tumor tissues. Joint analysis of eight spatial transcriptomic sections, including seven cancer specimens and a lymph node, identified 192 transcriptional programs that jointly resolved distinct tissue architectures (Figs. 4A-4C). While most tumor-lineage programs were sample-specific (Figs. S4A-S4K), multiple programs associated with common biological activities were shared across different tumors (Figs. 4D-4I). These included cell-cycle programs corresponding to G1/S, S/G2, and G2/M phases (Fig. 4D), as well as MYC-target, hypoxia, and type I interferon-response programs (Fig. 4E). We also identified shared programs in stromal and vascular populations (Fig. 4F), myeloid cells (Fig. 4G), and lymphoid populations (Fig. 4H). Importantly, these shared programs often coexisted with tumor-specific programs within the same cells (Figs. S4L and S4M), demonstrating that molecular inference can distinguish tumor-specific activities from cellular activities shared across diverse cancer types.

**Figure 4.**
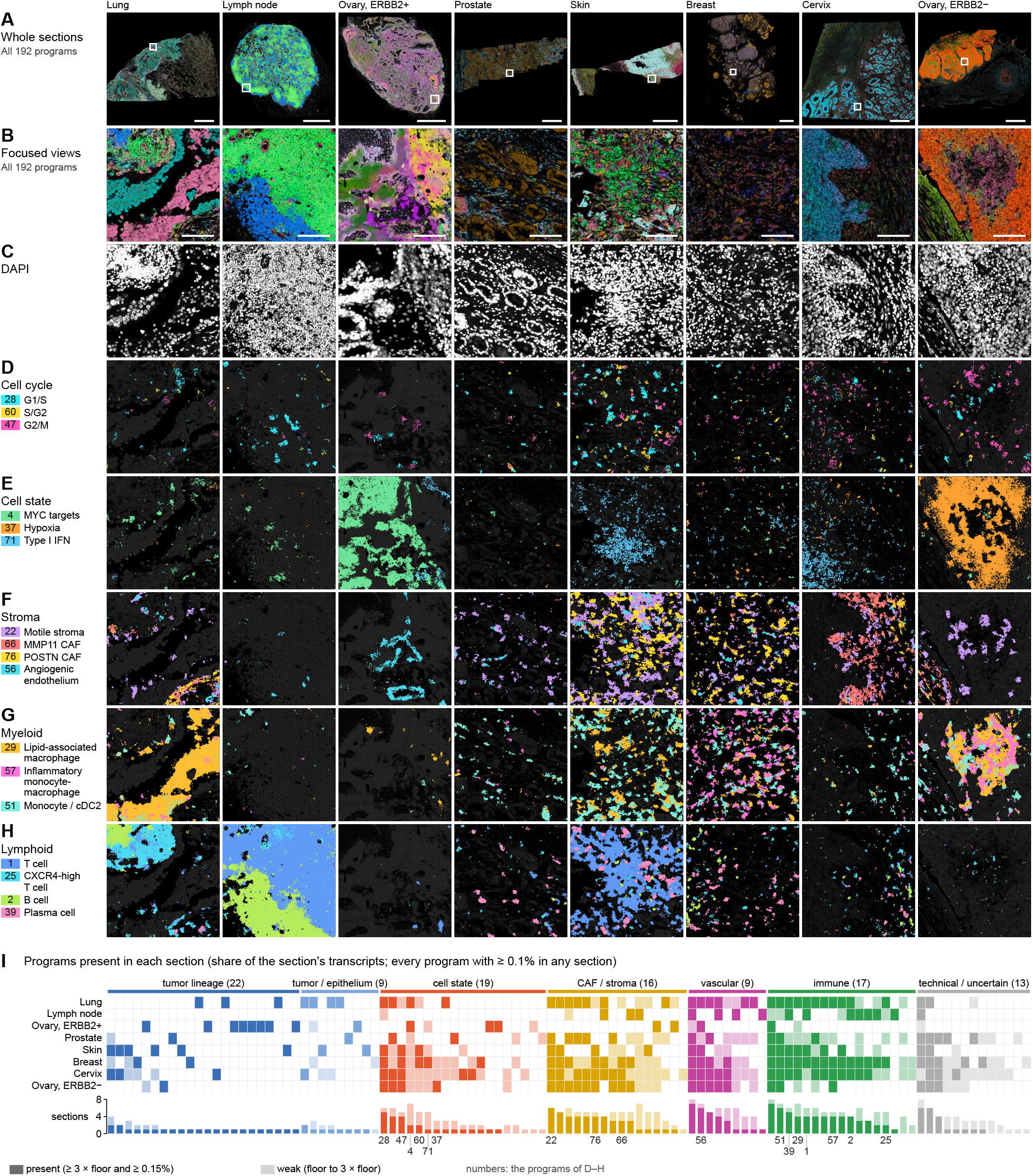
One FICTURE2 model fitted jointly to seven tumors and a lymph node resolves programs shared across cancer types. **(A)** Xenium Prime 5K sections of seven tumors and a lymph node, colored by the 192 programs of one jointly fitted model, so each program has the same color in every section; boxes mark the views in B-H. **(B)** Focused views of the boxed regions in A. **(C)** DAPI image of the views in B. **(D)** Cell-cycle programs in the views of B, detected in sections of different cancer types. **(E)** Cell-state programs, as in D. IFN, interferon. **(F)** Stromal and vascular programs, as in D. CAF, cancer-associated fibroblast. **(G)** Myeloid programs, as in D. cDC2, conventional type 2 dendritic cell. **(H)** Lymphoid programs, as in D. **(I)** Presence of each program in each section, called against a detection floor estimated from tumor-lineage programs in the sections of other tumors. Bars count the sections in which a program is present (dark) or weak (light). Most tumor-lineage programs occur in one section; cell-state, stromal, vascular and immune programs recur.

### Reference-guided molecular inference across multiple single cell and spatial transcriptomic technologies

FICTURE2 could also transfer cell types and transcriptional programs defined in single-cell data to spatial datasets generated by different technologies. Using a colorectal cancer scRNA-seq study^6^ with both single cell and ST experiments, we projected 39 published cell-types and 96 transcriptional programs learned by latent Dirichlet allocation (LDA) from the scRNA-seq dataset onto Xenium and Visium HD datasets from serial sections of three tumor specimens (P1, P2, and P5).^6^ Despite the distinctions between imaging- and sequencing-based technologies including having different gene sets (422 vs 18,055 genes), cell-type projections on data from the two technologies produced highly concordant tissue architectures (Figs. 5A-5C) and consistent global cell-type composition (Spearman ρ = 0.87–0.88; Fig. S5A) across all three specimens. FICTURE2 recovered fine-scale distributions of malignant epithelial, stromal, and immune populations in corresponding regions (Figs. 5D-5F), while such projections were reported unsuccessful in the original study.

**Figure 5.**
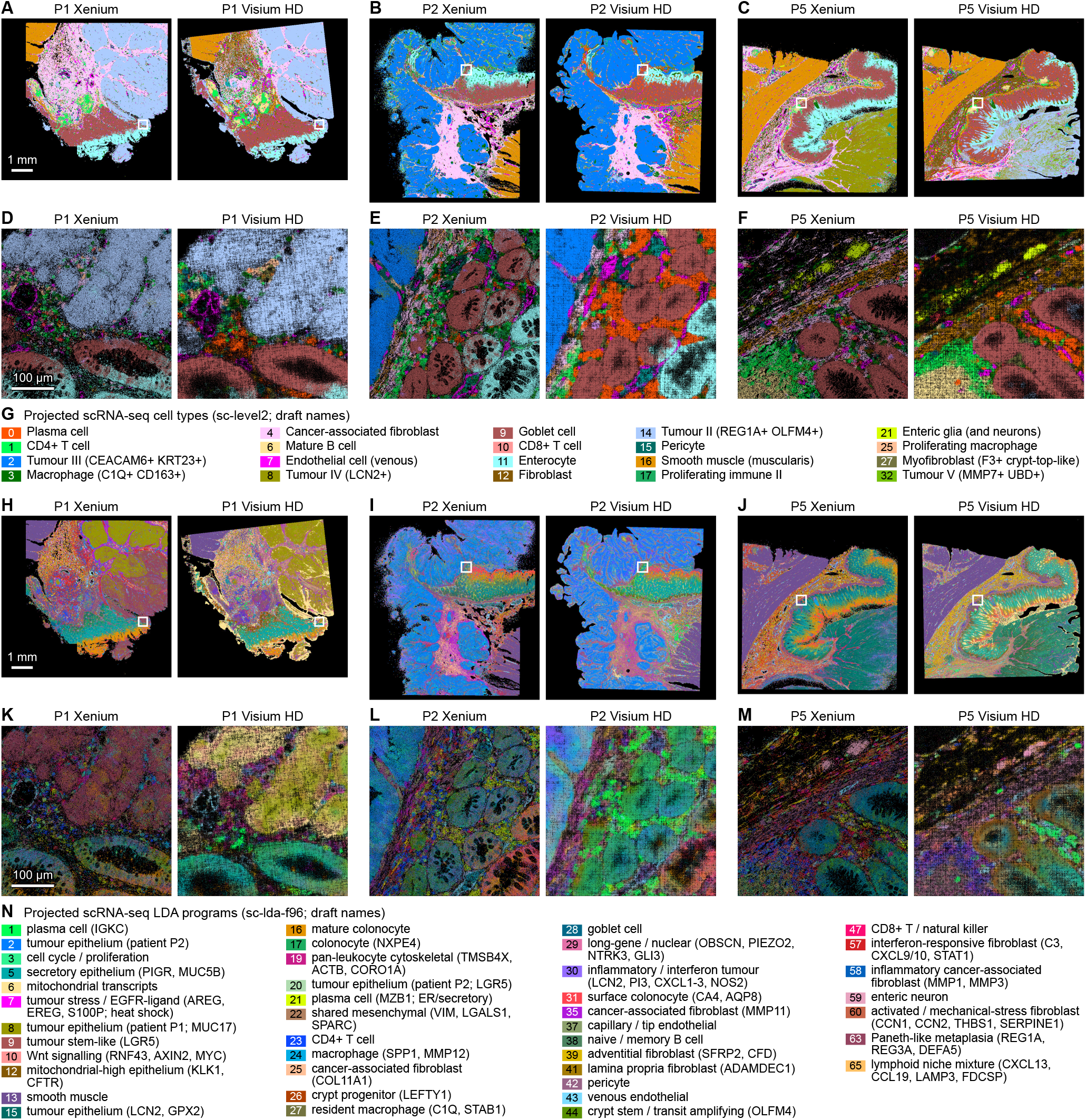
Projected scRNA-seq cell types agree across Xenium and Visium HD colorectal cancer sections more closely than projected programs. **(A)** Projected scRNA-seq cell types (key in G) on serial sections of colorectal cancer specimen P1 profiled by Xenium (left) and Visium HD (right). Box, region enlarged in D. **(B)** As in A, for P2. Box, region enlarged in E. **(C)** As in A, for P5. Box, region enlarged in F. **(D)** Enlargement of the box in A. Compartments occupy corresponding positions on the two platforms, but individual glands and cells are not matched one to one because the sections are serial. **(E)** As in D, for the box in B. **(F)** As in D, for the box in C. **(G)** Cell-type key for A–F. Tumor types are patient specific (Tumor II, P1; Tumor III, P2; Tumor IV, P5; Tumor V, a patient without spatial data). **(H)** The sections of A colored by the 96 LDA programs learned from the scRNA-seq reference and projected onto the molecules (key in N). Box, region enlarged in K. **(I)** As in H, for P2. Box, region enlarged in L. **(J)** As in H, for P5. Box, region enlarged in M. **(K)** The region of D colored by projected programs. Major compartments remain recognizable, but local program composition differs more between platforms than cell-type composition does in D. **(L)** As in K, for the region of E. **(M)** As in K, for the region of F. **(N)** Program key for H–M; genes in parentheses are representative program genes; program colors are unrelated to the cell-type colors in G.

In contrast, projection of the 96 scRNA-seq-derived transcriptional programs revealed substantial differences between Xenium and Visium HD (Figs. 5H–5J). Although major spatial patterns remained comparable, we observed lower cross-platform concordance in global program abundance (Spearman ρ = 0.55–0.65; Fig. S5A) and local program distributions (Figs. 5K–5M). Notably, the representation of each program’s genes in the targeted Xenium panel drives these differences (Fig. S5B). For example, the plasma cell type (*IGKC*, *JCHAIN*), mesenchymal program (*VIM*, *LGALS1*, *SPARC*), and mitochondrial program were abundant in Visium HD but scarce in Xenium, whose targeted panel lacks their defining genes (Fig. S5B). Nevertheless, program-level spatial agreement is compelling (Fig. S5C) and programs projected from scRNA-seq showed correspondence with programs independently inferred from each spatial dataset (Figs. S5D and S5E). Thus, FICTURE2 enables reference-guided inference of both cell types and transcriptional programs across distinct spatial technologies, which then demonstrates that fine-grained program inference fully realizes the advantage of whole transcriptome measurements that is undervalued if analyzed only through cell-type annotations.

### Segmentation-free molecular inference reveals disease-associated expression changes beyond cellular composition

Previous studies have shown that cell segmentation errors confound differential expression tests.^21^ FICTURE2 provides a natural alternative to identify disease-associated changes in both transcriptional program composition and individual gene expression within each program. We analyzed a published idiopathic pulmonary fibrosis (IPF) lung dataset from Banovich and colleagues,^22^ comprising 25 Xenium tissue cores from controls (n = 7) and regions with less (IPF-less, n = 7) or more extensive fibrosis (IPF-more, n = 11; Fig. 6A). FICTURE2 resolved fine-scale cell-type distributions directly from individual molecules (Figs. 6B and 6C), bypassing the confounding from segmentation errors. IPF-more regions exhibited extensive changes in cell-type and transcriptional program composition (Figs. 6D and 6E; Figs. S6A and S6B), with distinct fibroblast programs separating disease-related and unaffected activities and revealing more interpretable heterogeneity than conventional cell-type classifications (Fig. S6C).

**Figure 6.**
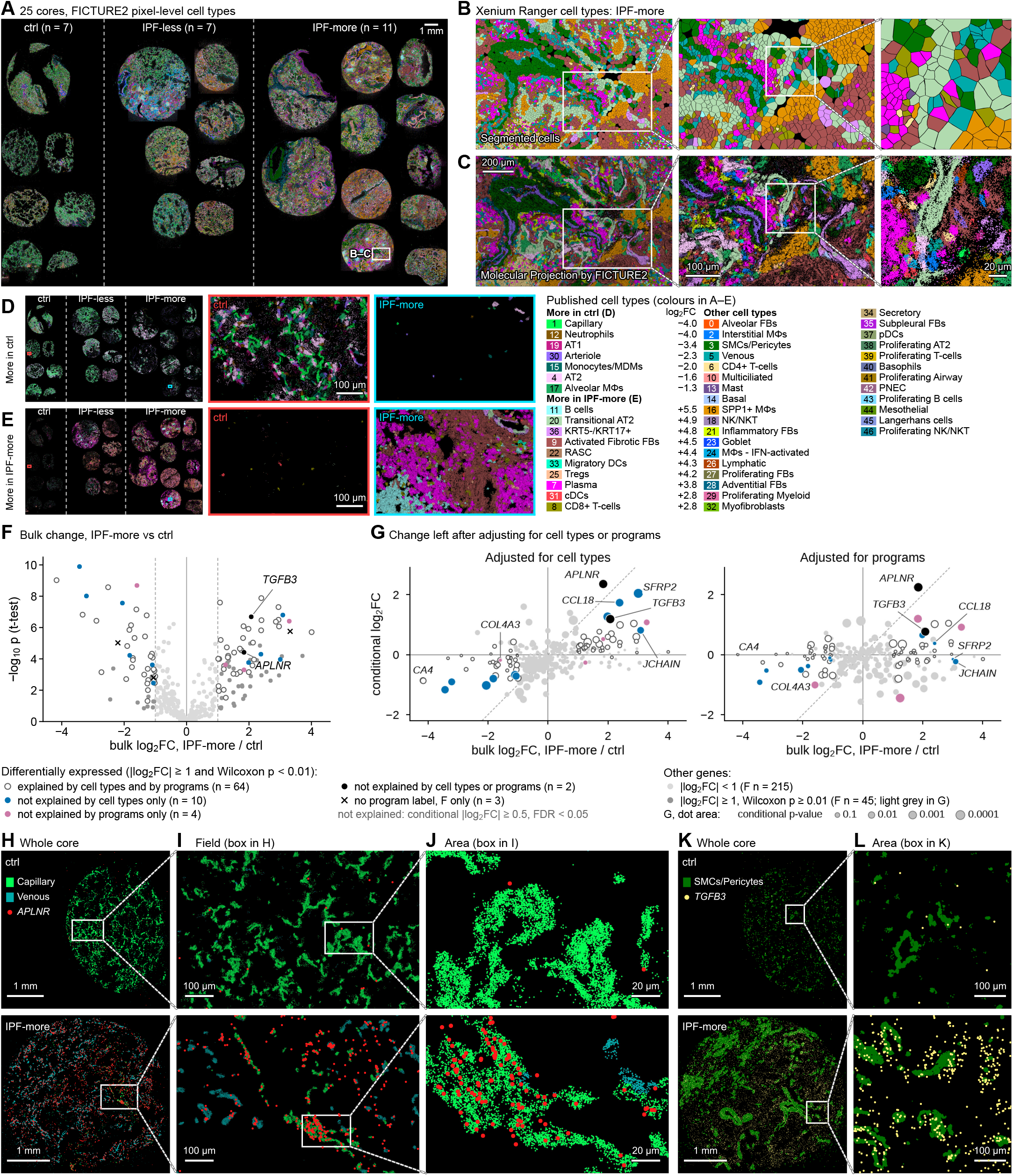
Disease-associated expression changes in IPF lung accounting for cell-type and program composition. **(A)** Published cell types projected by FICTURE2 onto all 25 lung cores from control, less-affected (IPF-less) and more-affected (IPF-more) lung; the box marks the region enlarged in B and C. **(B)** The region boxed in A at successive magnifications, showing Xenium Ranger-segmented cells colored by their published cell type. **(C)** The same fields as in B, with each molecule colored by the cell type assigned by FICTURE2 projection of the published cell-type profiles, without cell boundaries. **(D)** Cell types more abundant in ctrl than in IPF-more cores (FDR < 0.05, two-sided Wilcoxon rank-sum test on per-core shares), shown alone on all cores (left) and in the boxed control and IPF-more regions (right); log₂FC (key), ratio of molecule shares pooled per group. **(E)** As in D, for cell types more abundant in IPF-more than in control cores. **(F)** Bulk expression change of the panel genes between IPF-more and control cores; p, t-test with cores as units. Differentially expressed genes are marked by whether their change is explained by cell-type or program abundance in G. **(G)** Change left after adjusting each core’s expected expression for the abundance of published cell types (left) or FICTURE2 programs (right), against the bulk change in F, for the 319 genes labelled in both layers. Dot area, permutation p-value; symbols as in F. **(H)** All *APLNR* transcripts, with the molecules of the Capillary and Venous cell types, in a whole control (top) and IPF-more (bottom) core; other molecules are not shown. **(I)** Fields boxed in H. **(J)** Areas boxed in I; in IPF-more lung, *APLNR* transcripts lie within the capillary signal. **(K)** As in H, for all *TGFB3* transcripts with the molecules of the SMCs/Pericytes cell type. **(L)** Areas boxed in K.

These changes in cell-type and program compositions explain most of the differential gene expression between IPF-more and control tissues, which can be explained by conditional analysis on cell-type or transcriptional program abundance (Figs. 6F and 6G; Fig. S6D). However, 10 genes exhibited changes unexplained by cell-type composition alone, four by program composition alone, and two by either (Fig. 6F). These residual changes indicate expression shifts of specific genes within existing cell types or programs, rather than changes attributable solely to their relative abundance.

*APLNR* and *TGFB3*, the two genes explained by neither cell types nor programs, highlighted potentially important disease-associated responses (Fig. 6G). *APLNR*, encoding the apelin receptor implicated in endothelial homeostasis and vascular remodeling,^23^ was markedly upregulated within capillary and venous endothelial populations (Figs. 6H–6J and S6E). *TGFB3*, an established driver of fibrotic disease,^24^ was strongly upregulated within smooth muscle/pericyte populations (Figs. 6K and 6L; Fig. S6E), identifying a perivascular source of this profibrotic factor. Thus, segmentation-free molecular inference can pinpoint disease-associated gene regulation within specific cell populations, revealing candidate disease drivers and signaling pathways that would otherwise be obscured by extensive changes in tissue composition.

### FICTURE2 combines molecular precision, biological interpretability, and computational scalability

Finally, we benchmarked FICTURE2 against existing spatial annotation methods to evaluate whether single-molecule inference could improve biological resolution without compromising computational efficiency. In a developing mouse tissue section, Space Ranger annotations based on 16-μm and 8-μm bins or segmented cells, FICTURE1’s pixel-level inference, and FICTURE2’s molecule-level inference all reproduced major anatomical structures at the whole-section scale (Figs. 7A–7F). However, striking differences emerged at higher magnification (Figs. 7G–7L). In cartilage maturation regions, conventional bin- and cell-based annotations produced coarse or fragmented spatial patterns, while FICTURE1’s pixel-level inference provided finer but relatively noisy assignments. In contrast, FICTURE2’s molecule-level inference resolved sharply delineated, continuous transcriptional patterns corresponding to distinct cartilage maturation zones and surrounding tissue layers (Fig. 7L). Similar improvements were observed across osteoblast and perichondrial layers, mesenchymal and vascular structures, and localized cell-cycle programs (Figs. S7A–S7J), demonstrating that single-molecule inference resolves fine-scale biological organization obscured by spatial aggregation.

**Figure 7.**
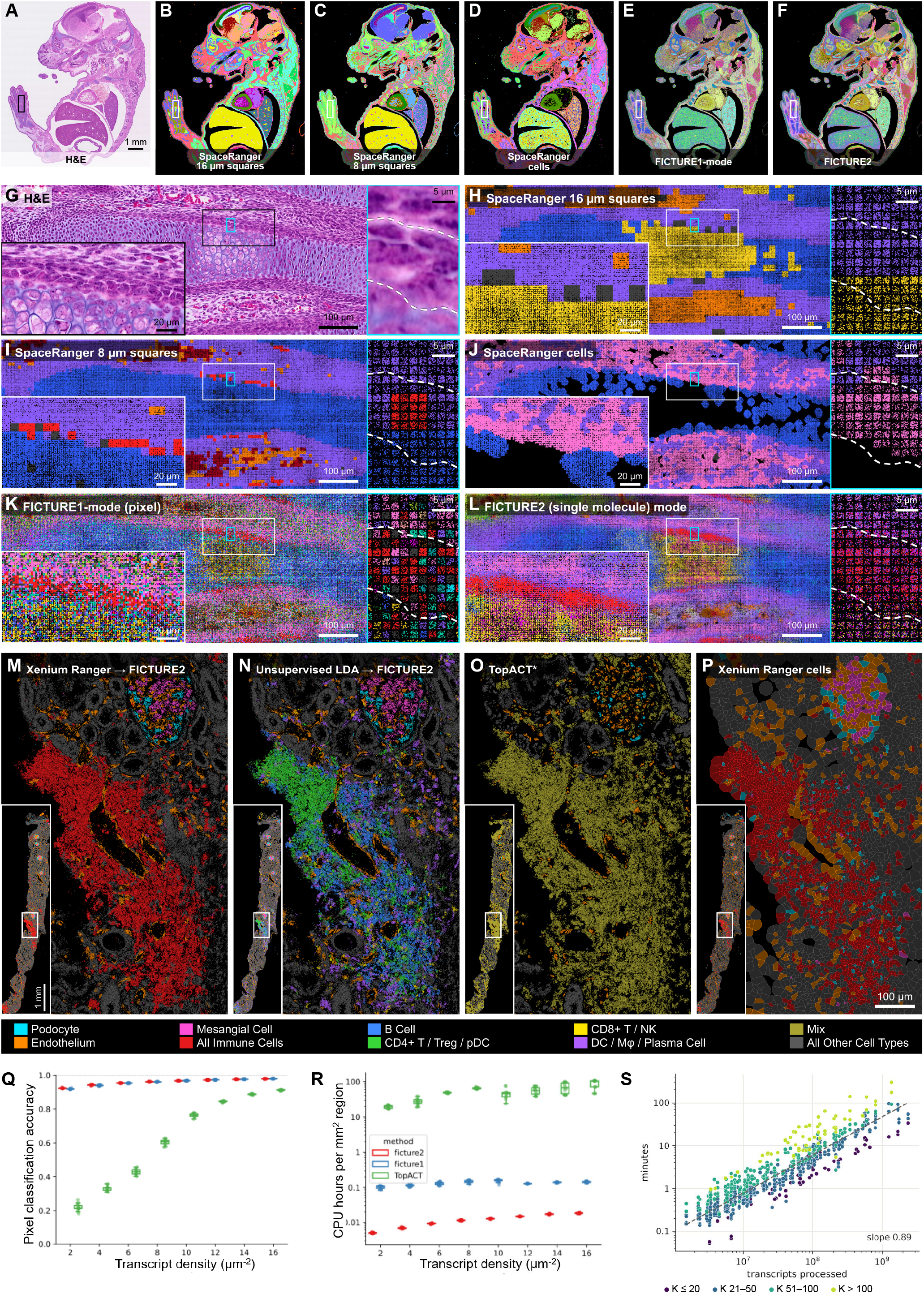
Comparison of single-molecule inference by FICTURE2 with FICTURE1 and other methods. **(A)** H&E image of a sagittal mouse embryo (E17) section profiled by Visium HD; box, the limb region enlarged in G–L (rotated 90°). **(B)** The section in A labelled by Space Ranger graph-based clusters of 16-µm square bins. Colors in B–D are not matched across methods; E and F share program colors. **(C)** As in B, for 8-µm square bins. **(D)** As in B, for Space Ranger cells segmented from the H&E image. **(E)** The section labelled by a 192-program LDA model decoded in FICTURE1 mode, which assigns one program to each 0.5-µm pixel. **(F)** The same model and transcripts decoded by FICTURE2, which assigns a program to each transcript. **(G)** H&E of the region boxed in A, a digit cartilage element. Lower left, the black-framed window enlarged; right, the cyan box enlarged; dashed line, the compartmental boundaries traced. **(H)** As in G, for Space Ranger 16-µm square bins; labels are colored to match FICTURE2 programs, grey if not in the key (key in Fig. S7). Transcripts are drawn at random offsets within their 2-µm bin. **(I)** As in H, for 8-µm square bins. (J) As in H, for Space Ranger cells. Transcripts outside segmented cells (black) made up 10.4% of the region, 91% of them within the chondrocyte labels of H. **(K)** As in H, for the 192-program model decoded in FICTURE1 mode; all transcripts of a 2-µm bin take one program (right). **(L)** As in K, decoded by FICTURE2; transcripts are assigned individually, so one bin can hold several programs (right). **(M)** A glomerulus (top) and an immune infiltrate in a Xenium section of human kidney with IgA nephropathy, with transcripts assigned by FICTURE2 to the cluster profiles of Xenium Ranger-segmented cells (FICTURE2-XR). Inset, whole section. **(N)** As in M, with transcripts assigned to 48 programs that LDA learned without segmentation (FICTURE2-LDA). **(O)** As in M, for the published TopACT annotation, whose “Other” class is shown as Mix. **(P)** As in M, for Xenium Ranger-segmented cells colored by their cluster. **(Q)** Pixel classification accuracy (fraction of pixels assigned their simulated cell type) of FICTURE2, FICTURE1 and TopACT on simulated tissue across transcript densities; boxes summarize replicate simulations (colors as in R). **(R)** Computational cost of the analyses in Q, in CPU hours per mm² of simulated tissue. **(S)** Wall-clock time of 829 FICTURE2 decodes of real data (each of one sample against one model with K factors, 24 threads) from 94 analyses on ten platforms, against transcripts processed; dashed line, least-squares fit on log–log axes.

We further compared FICTURE2 with TopACT,^25^ a recently developed method for subcellular spatial cell-type annotation, in complex kidney regions containing glomeruli and immune infiltrates. Both reference-guided and unsupervised FICTURE2 inference resolved podocytes, endothelial cells, mesangial cells, and distinct immune populations, whereas TopACT frequently assigned overlapping populations to a mixed category (Figs. 7M–7P). Marker-based evaluation confirmed greater molecular specificity: FICTURE2 assigned up to 89.3% of mesangial marker transcripts to the corresponding identity, whereas TopACT, which has no mesangial class, placed 39.1% in its mixed class (Figs. S7K and S7L). Within immune infiltrates, FICTURE2 distinguished B cells, T-cell subsets, plasma cells, and myeloid populations that TopACT largely classified as mixed (Figs. S7M–S7O). Moreover, molecule-level inference preserved immune marker specificity without requiring cell segmentation or arbitrary spatial binning (Fig. S7P).

These gains in interpretability were achieved without sacrificing accuracy or scalability. Across transcript densities, FICTURE2 maintained classification accuracy comparable to FICTURE1 and substantially higher than TopACT, particularly under sparse sampling, where accuracy remained approximately 94% at 2 transcripts/μm² compared with approximately 20% for TopACT (Fig. 7Q). Remarkably, FICTURE2 required approximately an order of magnitude less CPU time than FICTURE1 and thousands-fold less than TopACT (Fig. 7R). Its runtime scaled approximately linearly with transcript count, supporting analyses with billions of molecules (Fig. 7S). Together, these results establish FICTURE2 as a computationally scalable approach that combines high classification accuracy with the molecular resolution needed to decipher complex tissue organization and overlapping biological programs.

## Discussion

Spatial transcriptomics has advanced in sensitivity, molecular coverage, and spatial resolution, now routinely measuring millions to billions of individual RNA molecules within each intact tissue section. FICTURE2 opens new opportunities to investigate physiological activities at this molecular resolution and at scale. By inferring transcriptional programs and profiling their spatial distributions directly from individual transcripts, FICTURE2 can resolve cellular identities, differentiation states, metabolic activities, proliferation, and responses to environmental signals as coexisting programs, as well as examine how these activities are organized within cells, coordinated across tissue compartments, and remodeled during development and disease.

Our findings illustrate how transcriptional programs provide more continuous and multidimensional interpretations of cellular behavior. During epidermal wound healing, differentiation programs persisted as injury-response programs gradually emerged, revealing progressive molecular remodeling that was largely obscured by discrete cell-type assignments. Developing skeletal muscle similarly exhibited overlapping programs associated with myofiber maturation, fiber-type specification, and neuromuscular junctions. Across diverse cancers, shared cell-cycle, hypoxia, MYC-associated, and inflammatory programs coexisted with tumor-specific transcriptional identities. These observations suggest that cellular plasticity can be understood through the selective activation, persistence, and reorganization of transcriptional programs. Mapping these changes across developmental stages, disease progression, and therapeutic interventions could reveal common principles governing cellular adaptation across tissues.

A particularly striking discovery was the subcellular organization of transcriptional programs in colorectal cancer. Previous studies documented polarized mRNA localization in intestinal epithelia using subcellular microdissection and transcript imaging,^26,27^ while recent spatial transcriptomic studies have characterized intracellular RNA localization across tissues.^28^ FICTURE2 extended these observations to coordinated transcriptional programs, identifying distinct apical, basal, lateral, and nuclear programs through unsupervised molecular inference. These polarized programs coexisted with proliferation, metabolic, and inflammatory activities, exposing multiple dimensions of molecular heterogeneity within the same tumor. Such analyses provide opportunities to investigate how intracellular RNA organization relates to epithelial polarity, localized translation, malignant progression, and interactions with the surrounding microenvironment. More broadly, transcriptional programs may reveal spatially organized functional domains across tissues whose cellular architecture and molecular organization remain poorly characterized.

Molecular inference also provides a powerful approach to identifying disease-associated gene regulation within specific cell populations. In idiopathic pulmonary fibrosis, extensive cellular remodeling accounted for most changes in bulk gene expression, whereas composition-adjusted analysis identified genes whose expression increased disproportionately within particular cell populations. These included *APLNR* in capillary and venous endothelial populations and *TGFB3* in smooth muscle/pericyte populations. The apelin–APLNR signaling axis has important roles in vascular homeostasis and remodeling,^23^ and recent studies have implicated altered pulmonary endothelial *APLNR* expression in lung injury and fibrosis.^29^ TGFβ3 has established functional roles in fibrotic disease.^24^ Identifying these changes within their specific cellular contexts provides a more precise view of disease-associated signaling and nominates potential regulators for functional investigation. This approach can be extended to systematically identify disease-associated genes and programs whose regulation is obscured by extensive changes in tissue composition.

The growing role of transcriptional programs in connecting genes to phenotypes creates further opportunities for molecular spatial inference. Single-cell factorization has established the coexistence of cell-identity and activity programs,^16^ while large-scale Perturb-seq studies increasingly define how individual genes regulate coordinated biological programs. Recent work has connected these regulatory relationships to human genetic variation, disease susceptibility, and complex traits.^15,30^ FICTURE2 provides the spatial context for investigating these relationships in intact tissues. Programs defined through genetic perturbations can be projected onto spatial transcriptomes to identify the cellular populations and tissue environments in which their associated regulatory activities occur. Conversely, disease-associated programs discovered through spatial inference can guide the selection of perturbation targets and experimentally testable phenotypes. Integrating molecular spatial inference with genetic perturbations and human genetic studies could establish systematic connections among genetic regulators, transcriptional programs, their spatial organization, and disease phenotypes.

An equally important advance is the generality and computational efficiency of FICTURE2. Each spatial transcript contributes only two essential observations: its molecular identity and spatial coordinates. This minimal input makes the method applicable across diverse spatial transcriptomic technologies, including imaging- and sequencing-based platforms, using a common inference procedure. The shared molecular representation provides a foundation for harmonizing datasets generated by different technologies and experimental protocols. FICTURE2 can learn transcriptional programs directly from spatial data or project cell-type and transcriptional program profiles from existing single-cell references, allowing the same biological references to be applied across technologies. Our original FICTURE method already enabled segmentation-free pixel-level inference with orders-of-magnitude greater computational efficiency than previous approaches.^17^ FICTURE2 advances this capability to individual-molecule inference while reducing computational cost by another order of magnitude relative to FICTURE and by thousands-fold relative to TopACT in our benchmarks. Its near-linear scaling makes comprehensive analysis of massive spatial transcriptomic collections practical. Consequently, datasets can be systematically reprocessed as improved reference atlases become available, alternative transcriptional programs are developed, and new biological hypotheses emerge. These capabilities are further extended through CartoScope and CartoAgent, described in an accompanying study.^18^

Together, molecular-level inference, cross-platform compatibility, and computational scalability create the foundation for a new generation of functional tissue atlases. Large collections of spatial transcriptomes can be analyzed using shared molecular representations, allowing conserved biological programs to be compared across tissues, individuals, diseases, and technologies. Such atlases could reveal how common cellular activities are deployed in distinct anatomical contexts, how interacting programs organize tissue microenvironments, and how pathological programs emerge and spread during disease progression. As transcriptional references, perturbation datasets, and regulatory models continue to expand, existing spatial transcriptomic data can be repeatedly reinterpreted to address increasingly sophisticated biological questions. By resolving transcriptional programs at individual-molecule resolution across entire tissue collections, FICTURE2 enables systematic investigation of the spatial organization and regulation of biological function.

## Methods

### Molecule-level spatial factor inference

FICTURE2 is a complete reimplementation and extension of the spatial factor model of FICTURE,^17^ which we briefly summarize here. Let *β_k_* ∈ Δ*^M^*^−1^ be the compositional gene-expression profile over *M* genes for factor *k* ∈ [*K*], representing a transcriptional program. During spatial decoding the *K* profiles are held fixed by default. Spatial context is shared through anchors, lattice points denser than typical single cells. Each anchor *j* carries factor proportions *θ*_j_ ∈ Δ*^K^*^−1^, informed by observations within a neighborhood. An observation at location **y***_i_* contributes *ψ_i_*_j_ of its molecule counts to anchor *j*, *ψ_i_*_j_ > 0 only for anchors within a support radius *r*. Based on *ψ_i_*_⋅_ and *θ*_j_ for neighboring anchors, each observation then obtains its posterior factor probabilities *φ_i_* ∈ Δ*^K^*^−1^. The local latent variables *θ*, *φ*, and *ψ* are inferred by mean-field variational inference. FICTURE2 supports three inference modes, differing in what counts as an observation and how gene identity enters the likelihood.

#### Pixel mode (original FICTURE)

In the original pixel mode, molecules are first binned into square pixels of side length *s* (typically 0.5–1 µm). The observation *i* is a pixel with gene-count vector 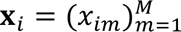. All molecules in a pixel share a single anchor assignment *c_i_* and a single factor assignment *z_i_*, and **x***_i_* ∣ *z_i_* = *k* ∼ Multinomial(∑*_m_ x_im_*, *β_k_*). The coordinate-ascent updates are

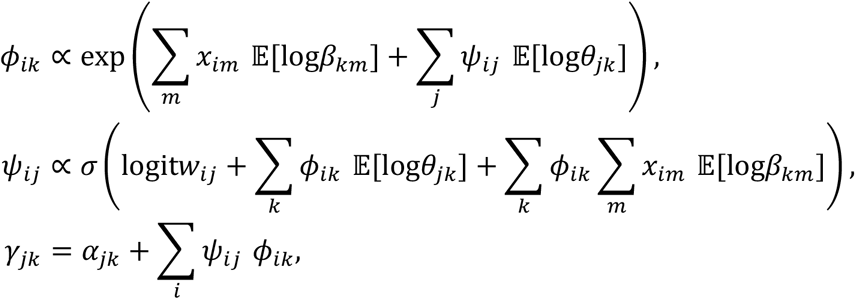

where *σ* is the logistic function and *ψ_i_* is normalized over the anchors within distance *r* of pixel *i*.^17^ Because the gene term sums over every gene in the pixel, all molecules in a pixel receive the same factor posterior *φ_i_*. This is a reasonable assumption when the factors are mutually exclusive cell types and lateral diffusion is negligible (as for most imaging-based ST platforms). It does not hold when the factors are transcriptional programs that can co-occur within a cell or within a single pixel.

#### Single-molecule mode

Every molecule is its own observation and is decoded at its recorded coordinates, with no spatial binning. Let molecule *i* have gene identity *g_i_*, location **y***_i_*. The generative process of FICTURE is applied to each molecule:

1. Draw the anchor assignment *c_i_* ∼ Categorical(**w***_i_*).
2. Draw the factor assignment *z_i_* ∼ Categorical(*θ_ci_*).
3. Draw the gene identity *g_i_* ∼ Categorical(*β_zi_*).

Two molecules from different genes at the same exact location can thus have different posterior factor probabilities when their genes are characteristic of different programs that are both active locally. Molecules of non-specific genes, by contrast, are attributed mainly according to the local composition. Single-molecule mode is suitable for imaging-based platforms (e.g., 10X Xenium, Atera, MERFISH, CosMx SMI) and submicron-resolution sequencing-based platforms when the observation is sparse.

#### Feature-pixel combination mode

This mode collapses molecules of the same transcripts within the same spatial location, allowing different transcripts to be assigned to different transcriptional programs just like single-molecule inference without overly sacrificing computational efficiency. This is particularly useful for sequencing-based platforms with subcellular but not optical resolution, such as 10X Visium HD (2*μm* pixels) and Illumina StrataMap (1*μm* pixels), where it is common to observe multiple molecules of the same gene within a single pixel.

### Factor-conditional differential expression

To compare gene expression between conditions within a given cell type or program, we used a factor-conditional differential expression (DE) test. The test uses the pixel or molecule level probabilistic factor assignments directly, without using cell segmentation.

#### Soft, factor-stratified aggregation

We divide each tissue sample into square spatial units *u* of side length *L* (e.g., 20 µm). Each molecule contributes to every factor *k* for which its posterior probability *φ_ik_* is at least *p*_min_ (default 0.01). The soft factor-specific count of gene *m* in unit *u* and the corresponding unit total are

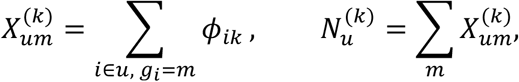

where *φ_ik_* is the posterior probability of molecule *i* belonging to factor *k*; 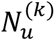 is the factor-stratified count, summed over the tested gene panel. A unit is included in the analysis of factor *k* only if 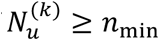 (default 10).

For simplicity, we consider comparing two groups, where each pixel is assigned to at most one group. In practice, the contrast can be between samples sharing the same set of factors, or between regions within a sample.

#### Test statistic

For each factor *k* and gene *m*, we model the counts as

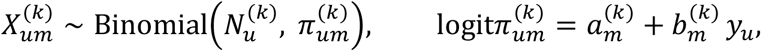

where *y_u_* ∈ {0,1} indicates the group of unit *u*. The null hypothesis is 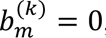, that the relative expression of gene *m* among molecules attributed to factor *k* does not differ between groups. This design has a closed-form maximum-likelihood estimate. With U_g_ the set of units in group 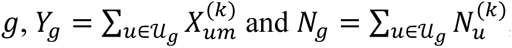, the estimates are

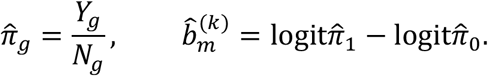

To stabilize estimates for genes that are rarely or never observed in one group, each 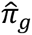 is bounded below by 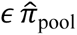. Here 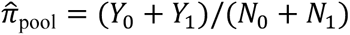 and *ε* = 0.05 by default.

Molecules within a spatial unit are not independent, and the expression of a gene varies across units more than binomial sampling predicts. To account for this, the standard error of *b̂* is computed with a heteroskedasticity-robust (sandwich) estimator that treats units as the sampling units:

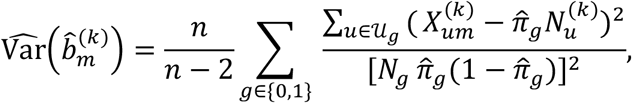

where *n* = |U_0_| + |U_1_| and the factor *n*/(*n* − 2) is the HC1 small-sample correction. Two-sided p-values are obtained from the Wald statistic 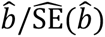 under a standard normal reference. Tests are performed only for (factor, gene) pairs with *Y*_0_ + *Y*_1_ greater than a threshold, 100 by default. For each retained (factor, gene) pair we report the log odds ratio *b̂*, the proportions *π̂*_0_ and *π̂*_1_, the total soft count and the p-value.

As an optional nonparametric calibration, units within factor *k* are permuted between the two groups *B* times while keeping the number of units per group fixed. The permutation p-value is the fraction of permutations with 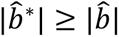.

#### Accounting for uncertain factor assignments

Because factor assignments are probabilistic and imperfect, we adjust *π*_g_ with an approximate deconvolution. To estimate the extent of convolution, we compute a factor-by-factor co-assignment matrix for each input, *S_kl_* = ∑*_i_ φ_ik_ φ_il_*. These confusion matrices are calculated within each group and row-normalized to give matrices **C**_0_ and **C**_1_.

For each gene, we then model the observed per-factor proportions as mixtures of latent factor-specific proportions:

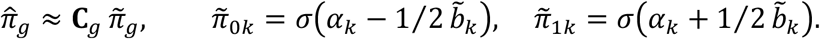

The parameters are estimated by minimizing the weighted least-squares objective

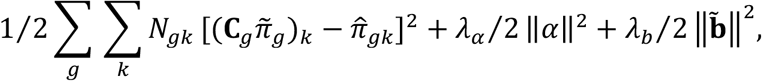

where *N*_g*k*_ is the total soft count of factor *k* in group *g*, *λ_α_* = 10^−6^ and *λ_b_* = 10^−2^. Minimization uses damped Gauss–Newton (Levenberg–Marquardt) iterations.

### Datasets

All datasets were downloaded from publicly available source (see **Data Availability**). Control or negative probes, unassigned codewords and, mitochondrial genes, and other non-gene features were removed prior to analysis. Visium HD counts were recorded per gene and 2-µm bin at the bin center; positions were jittered by up to ±0.8 µm in both directions for more informative visualization.

### Transcriptional program inference

Programs were learned by latent Dirichlet allocation (LDA) on transcripts aggregated in hexagons of 12 µm (for spatial transcriptomic datasets) or by cells (for scRNA-seq datasets). Every transcript was then assigned to programs by FICTURE2. A model fitted jointly to several sections gives each program the same identity in every section. Programs were named after their top marker genes, their location in the tissue and, where informative, the cell types that carry most of their transcripts. Names are interpretive labels; program numbers in the figures are the model’s factor indices.

### Wound-site analysis of mouse skin

#### Wound zones

At each of seven wound sites, an epidermal axis was traced by hand from the unwounded flank to the wound tip, and transcripts within 200 µm of it were binned in 20-µm steps; wound 01 (Fig. 1E,F) used a straight 1,240 × 174 µm strip. Seven epidermal cell types and eight epidermal programs were grouped by their distribution between unwounded and wounded epidermis (unwounded, shared and wounded cell types; unwounded, mild wound and severe wound programs; Fig. 1E,F), and group shares were computed among the transcripts of these labels only, excluding bins with fewer than 250 epidermal-program transcripts (800 for the wound 01 strip). Z2 began at the first bin where mild wound programs reached 20%, Z3 where severe wound programs reached 10%, and Z4 where they reached 50%; the preceding region was Z1. Program differences between zones are therefore partly expected by construction. Group shares in adjacent zones were compared by two-sided paired t-tests across sites (n = 7; uncorrected). The sites lie on one tissue microarray and may not come from independent animals.

#### Expression change explained by composition

For adjacent zones, each gene’s bulk log2 fold change (*x*) was compared with the change predicted from label composition alone (*y*), with each label’s expression profile held fixed and pooled over the two zones; zone pseudobulks were summed over sites. Genes with ≥10 transcripts in each of the two zones were included. The share explained was 1 − Z_g_*n*_g_(*x*_g_ − *y*_g_)²/Z_g_*n*_g_*x*_g_², with *n*_g_ the gene’s transcript count, computed for all sites together and per site.

#### Programs within a cell type

For cell type 25 (wound cornifying), the expected share of gene *g* in zone *z* was Z*_k_w*(*k*|*z*)*φ*(*g*, *k*), where *w*(*k*|*z*) is the posterior-weighted share of the cell type’s transcripts assigned to program k (sites pooled) and *φ*(·, *k*) the profile of transcriptional program *k* pooled over Z1–Z4.

### Facial muscle analysis of the neonatal mouse

We analyzed a 11.45 × 23.81 mm^2^ region of the day 1 mouse pup section containing 523.5 million transcripts with three inference layers: Xenium Ranger clusters of segmented cells, in which each transcript takes the cluster of its cell; the same cluster profiles assigned to individual transcripts by FICTURE2; and the 192 FICTURE2 programs. We focused on the analysis of 1.51 × 2.74 mm^2^ field of the head of the pup section containing facial muscle region. For the cross-tabulation of clusters and programs, the joint mass of each pair was the sum over transcripts of the product of the two posteriors. Enrichment was *log*_2_[(observed + 0.5)/(expected + 0.5)], with the expected mass taken under independence of the two layers. Tables built from the most probable label of each transcript, or from the clusters of segmented cells, gave the same five largest programs within the skeletal-myofiber cluster. Marker genes were tested for each label against all other labels of the same method within the region (chi-square test; fold change ≥ 1.5, P ≤ 0.001). The displayed genes combined known markers with the top three markers of each displayed label.

### Subcellular and microenvironment analysis of colorectal cancer

For the analysis of Xenium Prime 5K colorectal cancer sample from the SPATCH study,^20^ we used a 48-program LDA model and the published cell types of the segmented cells.

#### Depth zones

Tumor epithelium was divided by distance from the luminal surface into surface (0–0.5 mm), intermediate (0.5–2 mm) and deep (≥ 2 mm) zones; normal mucosa was kept separate.

#### Stroma-to-lumen axis

In a field of tumor glands, stroma was the area (1µm grid) where at least half of the smoothed transcripts carried a non-epithelial published cell type projected onto transcripts; stroma patches below 500 µm² were ignored. Gland midlines were the pruned medial axis of the non-stroma region. The relative distance of a transcript was *d_stroma_*/(*d_stroma_* + *d_midline_*), evaluated where the gland half-width was ≤ 75 µm. The masks used no programs or images. In published epithelial cells with ≥ 20 transcripts, a program was considered present when it was the top program of ≥ 3 transcripts; for each program pair, we compared the mean distance to stroma of their transcripts within cells carrying both.

#### Cell-cycle programs across lineages

Four of the 48 programs were identified as cell-cycle programs from their marker genes (3 S, 7 G1/S, 8 G2/M and 9 Mitotic; Fig. S3C); no predefined list of cell-cycle genes was used. For each published cell type (Epithelial cells split into tumor and normal; cells with at least 20 transcripts), we computed the share of its transcripts carried by each cell-cycle program (posterior mass). A cell was called cycling when the four cell-cycle programs together carried at least 20% of its transcripts. We report the fraction of cycling cells among all cells and among cells whose centroid lies at least 30 µm from the nearest epithelial cell centroid, to exclude spillover from tumor cells. To compare a program’s gene content between lineages, we built for each cell-cycle program and lineage group (epithelial; stromal: fibroblasts, endothelial cells, smooth muscle cells; immune) a profile over all 5,001 panel genes, from the transcripts assigned to the program inside cells of that group, and compared the profiles by cosine similarity.

#### Program coexistence

Programs were grouped into five classes (tumor core, cell cycling, lipogenic, neuroendocrine-like, inflammatory). In tumor epithelial cells with ≥ 50 transcripts, class presence was called with a primary rule and six alternatives (share ≥ 5% or ≥ 20%, ≥ 200 transcripts, top label, confident or nuclear transcripts only)

#### Microenvironment

Microenvironment programs were grouped by the region they border (deep tumor, tumor surface, normal mucosa). Across 250µm tiles, the abundance of each tumor program class was correlated (Spearman) with that of each microenvironment program within 50 µm outside the tumor. An association was considered to persist when the 95% block-bootstrap confidence interval (65 spatial blocks) of the correlation adjusted for depth and x, y position excluded 0.

### MAP7 immunohistochemistry images

Published MAP7 immunohistochemistry images from independent colorectal cancer specimens were obtained from antibody suppliers and the Human Protein Atlas to compare protein localization with FICTURE2 predictions (Fig. S3G).

### Joint analysis of eight tumor and lymph node sections

The eight sections (seven tumors and a reactive lymph node) were analyzed with the 192-program LDA model fitted jointly to all of them. The two ovarian sections are labelled by the *ERBB2* status of their tumor programs.

#### Program presence

The share of a program in a section was the fraction of the section’s transcripts assigned to it. The 105 programs with a share of at least 0.1% in at least one section were grouped into seven categories (Fig. 4I). Because a program carries posterior probabilities to some transcripts even where it is absent, we estimated a detection floor as the 95th percentile of the shares of tumor-lineage programs in the sections of other tumors, where their lineage cannot occur. A program was called present in a section when its share was at least three times the floor and at least 0.15%, and weakly present when its share lay between the floor and three times the floor.

Section-specific programs were selected from an automated multi-sample annotation of marker genes, spatial distributions, and host cells, reviewed by the authors. Tumor-associated programs were considered section-specific when present in only one section’s tumor compartment under the above rule, allowing weak calls elsewhere. General activity programs (cell cycle, hypoxia, MYC targets, and interferon response) were excluded. Squamous programs 33 and 36 were assigned to cervical carcinoma rather than skin, where they marked normal epidermis. This yielded 22 tumor section-specific programs. In the tumor-free lymph node, three nontechnical programs (30, 58, and 63) were selected based on presence only in that section and at least tenfold greater abundance than in any other section.

#### Programs within single cells

Transcripts were aggregated using the cell segmentations provided by Xenium Ranger. For each cell with at least 50 transcripts, the posterior mass of a program was calculated by summing its posterior probabilities across the cell’s transcripts, considering the top three programs assigned to each transcript. A cell was considered to carry a program when its posterior mass was at least 5 transcript equivalents and 3% of the cell’s transcripts. The program with the largest posterior mass was designated the dominant program.

For each pair of a tumor-specific and a conserved program, we computed the percentage of cells dominated by the tumor-specific program that also carried the conserved program (Fig. S4L; only tumor-specific programs dominating at least 100 cells were included). In the lymph node, which contains no tumor, cells dominated by the B-cell or T-cell program took the place of tumor cells.

### Single cell reference-guided projection across Xenium and Visium HD

We analyzed serial sections of published tumor specimens P1, P2 and P5 profiled by both Xenium and Visium HD.^6^ For Visium HD, two additional normal specimens (P3N and P5N) were also analyzed. The 39 published cell-type (level 2) profiles from the scRNA-seq atlas of the same study (260,506 cells from five tumors and three normal adjacent tissues) were projected onto every section with FICTURE2. The 96-program LDA model was learned from the single cell raw count matrix using FICTURE2 in the same manner to the unsupervised LDA applied in other spatial datasets, except that hexagons are replaced to single cells. Tumor cell types are patient specific. The 96-program LDA models from each spatial platform (Xenium and Visium HD) were also generated and served for comparison.

For each specimen, the Visium HD section was mapped onto the Xenium section by a rigid transform (rotation and translation) that maximized the similarity of rasterized maps of six projected cell-type groups and total tissue.

Label abundance was the sum over transcripts (UMIs on Visium HD) of the posterior probability of the label, expressed as a share of the section. Cross-platform agreement was summarized by the Spearman correlation of shares, and related to panel coverage, the fraction of each reference profile on the 407 Xenium genes, by Spearman correlation with the *log*_2_(VisiumHD/Xenium) share ratio. Spatial agreement was the Pearson correlation of each label’s share between platforms over shared tissue bins (100∼800 µm), averaged over labels with ≥ 0.5% of the mass, weighted by abundance. Projected scRNA-seq programs and each platform’s de novo programs, which decode the same molecules, were cross-tabulated per molecule within each section and pooled over specimens; enrichment was the log2 ratio of observed to expected joint mass under independence.

### Composition-adjusted differential expression in IPF lung

All tests compared the 7 controls/unaffected cores with the 11 IPF-more cores as independent replicates; IPF-less cores are shown for reference only. Exact tests enumerated all 31,824 assignments of group labels to the 18 cores, and FDR was controlled by the Benjamini– Hochberg procedure.

The abundance of a cell type or program in a core was its posterior-weighted share of the core’s molecules, compared by an exact two-sided Wilcoxon rank-sum test. Effect sizes are Hodges– Lehmann shifts in log₂ share with exact 95% confidence intervals (Fig. S6A,B) or log₂ ratios of shares pooled per group (Fig. 6D,E).

The bulk log₂FC of a gene was the log₂ ratio of its share of all molecules pooled over IPF-more and over ctrl cores; volcano p-values came from a t-test with core-clustered standard errors. Genes with |log₂FC| ≥ 1 and Wilcoxon p < 0.01 on per-core shares were considered differentially expressed.

To adjust for composition, each molecule took its top label in one layer (47 cell types or 48 programs). The expected count of gene *g* in core *c* was *E*_g*c*_ = Z*_k_N_kc_r*_g*k*_, with *N_kc_* the molecules of label *k* in *c* and *r*_g*k*_ the rate of *g* in *k* estimated from the other 17 cores. The conditional log₂FC was the IPF-more minus control mean of *log*_2_ ((*Y*_g*c*_ + 0.5)/(*E*_g*c*_ + 0.5)), with *Y*_g*c*_ the observed count, tested by exact permutation for the 319 genes labelled in both layers. A differentially expressed gene was not explained by a layer when its conditional |log₂FC| was ≥ 0.5 with FDR < 0.05. The fraction explained was 1 − (conditional/bulk log₂FC)²; in Fig. S6D up to three labels were added greedily to the model to show which account for it.

For *APLNR* and *TGFB3*, rates per 1,000 transcripts of segmented cells of a type (cores with ≥ 20 such cells) or per 1,000 molecules of a program (≥ 1,000 molecules) were compared by Wilcoxon test. To test whether a target label rose beyond the rest of the tissue, its rate was divided by the mean rate of comparison labels fixed from control data (the five most abundant labels outside the stromal–vascular compartment that were present in all cores and had a control rate < 25% of the target’s), and the change in mean log₂ ratio was tested by permutation.

For Fig. S6C, molecules of the control and IPF-more cores were pooled and counted by their top label, and marker genes were tested for each label against all other labels of the same layer (chi-square test; fold change ≥ 1.5, P ≤ 0.001).

### Benchmarking against binning, segmentation and TopACT

#### Visium HD mouse embryo

Space Ranger clusters of 16-µm squares, 8-µm squares and H&E-segmented cells were compared with the 192-program model trained by LDA, decoded in FICTURE1 mode or by FICTURE2. FICTURE1 mode decoded programs on a 0.5-µm pixel grid, each transcript taking the program of its pixel; as Visium HD places all transcripts of a 2-µm bin at its center, each bin received one program, so FICTURE1 mode effectively assign the same posterior for all transcripts within the 2-µm bin. FICTURE2 assigned each transcript on its own. The enlarged region was a 1,146 × 372 µm box around one digit cartilage element. There, each Space Ranger label took the color of the FICTURE2 program making up most of it, or else of the program it was most enriched for; labels outside the key were grey. A marker share was the fraction of a marker set’s transcripts in a region whose most probable label was the label of interest (here, in the box: *Col10a1*, hypertrophic chondrocyte program; *Top2a*, *Mki67* and *Cdk1* pooled, mitosis program).

#### Kidney comparison with TopACT

On a Xenium section of human kidney with IgA nephropathy,^25^ FICTURE2 assigned transcripts to the profiles of 24 Xenium Ranger cell clusters (FICTURE2-XR) or to a 48-program model (FICTURE2-LDA); both were compared with the published TopACT annotation, with labels grouped into shared cell types by marker genes. The largest immune infiltrate and the nearest glomerulus were detected from label density. As a proxy for ground truth, marker shares were computed for the marker sets in Fig. S7L,N against the matching label group, pooled over five glomeruli or over the infiltrate. As an upper bound for annotations with one label per unit, each Xenium Ranger cell and each square bin (2∼12 µm) in the infiltrate took the immune class contributing most of its marker transcripts.

#### Accuracy and runtime

FICTURE2, FICTURE1 and TopACT were run on tissue simulated as in the FICTURE evaluation^17^ at eight densities from 2 to 16 transcripts per µm². Accuracy was the fraction of pixels assigned their simulated cell type; cost was CPU hours per mm². Runtime on real data was the wall-clock time of 829 FICTURE2 decodes for the datasets deployed to CartoScope,^18^ each of one sample against one model (K = 10∼384), from 94 analyses on ten platforms with 24 threads, some running concurrently. Scaling was the least-squares slope of log time on log transcript count. All comparisons were descriptive.

## Code availability

The FICTURE2 software tool is publicly available at https://github.com/Yichen-Si/punkst and documented at https://yichen-si.github.io/punkst/. The scripts to reproduce the analysis in this manuscript will be available in a separate GitHub repository upon publication.

## Data availability

The Xenium wounded mouse skin data was downloaded from Gene Expression Omnibus (GEO) with accession ID GSE278611.^19^ The neonatal day 1 mouse pup data was downloaded from 10x Genomics public dataset collection at https://www.10xgenomics.com/datasets. The Xenium Prime 5K colon cancer dataset was downloaded from the SPATCH^20^ web site at http://spatch.pku-genomics.org/. Xenium Prime 5K multi-sample cancer datasets across six tumor types and lymph nodes are downloaded at 10x Genomics public dataset collection. For multi-sample CRC data, Visium HD, Xenium, and scRNA-seq Flex data was accessed from Gene Expression Omnibus under accession number GSE280318. The 25 IPF and unaffected lung tissue microarray biopsy cores are obtained from GSE250346, focusing on the 25 of 45 de-arrayed samples with individual DAPI images and segmented cell boundaries available. The Visium HD (11mm) mouse E17 embryo dataset was downloaded from 10x Genomics public data collection, and the Xenium human IgA nephropathy kidney biopsy sample was obtained from GSE264334 and Zenodo repository with DOI:10.5281/zenodo.10950538. All the output data generated from this analysis will be publicly available in the CartoStore, an AWS Open Data Registry, at https://registry.opendata.aws/cartostore/.

## Acknowledgements

This work was supported by the Taubman Institute (to H.M.K. and J.H.L.), the Chan Zuckerberg Initiative (to H.M.K. and J.H.L.), the Glenn Foundation for Medical Research (to J.H.L.), the Eric and Wendy Schmidt Center at the Broad Institute of MIT and Harvard (to Y.S.), and the Henry Sewall Legacy Professorship (to J.H.L.). Additional support was provided by the National Institutes of Health (R01HG011031, R01GM166173, and HHSN268201800002I to H.M.K.; UH3CA268091 and R01DK133448 to J.H.L.; and R01AG079163 to M.K. and J.H.L.).

## Ethics declarations

J.H.L. is an inventor on intellectual property related to Seq-Scope and has a potential financial interest through the University of Michigan in connection with Salus Biomed Co. H.M.K. has served as a paid consultant for Regeneron Pharmaceuticals. The remaining authors declare no competing interests.

## Declaration of generative AI and AI-assisted technologies in manuscript preparation

During the preparation of this manuscript, the authors used OpenAI ChatGPT, Google Gemini, and Anthropic Claude to assist with literature review, manuscript drafting, organization, and editing. CartoAgent, an AI-assisted spatial omics analysis system developed by the authors and operated primarily within the Anthropic Claude Code environment, was used to assist data exploration, analysis, and figure preparation. The authors critically reviewed and verified all reported results, analyses, figures, and scientific interpretations, and take full responsibility for the accuracy, integrity, and conclusions of this work.

## Supplementary Figure Legends

**Figure S1. Gradual transcriptional program changes at seven wound sites (A)** The seven wound sites of the tissue microarray (outlined, left) and each site along its epidermal axis, unwounded side on the left, colored by epidermal cell-type cluster (middle) and program (right). Wound 01 is the site in Fig. 1E,F. **(B)** Share of each cell-type (left) and program (right) group (as in Fig. 1E,F) in zones Z1–Z4 at the seven wound sites; two-sided paired t-tests between adjacent zones, uncorrected for multiple testing. **(C)** Marker genes of the epidermal programs (top) and cell-type clusters (bottom) across the whole tissue microarray. **(D)** Program composition within cell type 25 Wound cornifying across zones Z1–Z4; programs below 4% in every zone are pooled as other programs. **(E)** *Krt1* and *Arg1* within cell type 25 Wound cornifying: observed share and share expected from its program composition in D with program profiles held fixed (Methods).

**Figure S2. Molecule-level program analysis of neonatal mouse facial muscle. (A)** Marker genes of the five largest clusters (as in Fig. 2B) and eleven programs in the facial muscle region **(B)** Flow of facial muscle transcripts from segmented-cell clusters to the same clusters assigned per transcript (FICTURE2-XR; Fig. 2B) to programs (FICTURE2-LDA). All columns share one mass scale (percentages of all the region’s transcripts), so a layer that labels fewer transcripts is shorter; labels below 1% are pooled.

**Figure S3. Colorectal cancer programs vary with depth, layer within cells, coexist in single cells, span lineages and track the microenvironment. (A)** The published annotation labels all malignant glands Epithelial, whereas FICTURE2 programs vary within them (window, box at left). Bars, program composition of transcripts in published Epithelial cells per depth zone (defined in B). **(B)** Tumor programs by depth from the luminal surface: depth zones of the tumor epithelium (left), each program’s share of tumor epithelial transcripts along depth (middle) and the log2 ratio of deep to surface shares (right). **(C)** Marker genes of the epithelial programs across the whole section, with published Epithelial and proliferating T (Tprolif) cells for comparison; dot size and color are computed within each method. **(D)** Distance of tumor epithelium to the main stroma in a field of glands; gland midlines mark the lumen end of the axis in F. **(E)** Distance to the main stroma of the transcripts of each subcellular program in the field of D. **(F)** Program fractions along the relative stroma-to-midline distance (left), their medians under different gland half-width limits (second), the percentage of each program’s transcripts inside segmented nuclei (third) and, among cells carrying both programs of a pair, the percentage in which the first lies closer to the stroma (right). **(G)** MAP7 immunohistochemistry showing preferential localization to the apical, luminal side of colorectal tumor glands. Images reproduced from Thermo Fisher Scientific (PA5-31782; © 2025 Thermo Fisher Scientific Inc. Used under permission; www.thermofisher.com), Fortis Life Sciences/Bethyl Laboratories (A301-490A and A301-534A; reproduced with written permission), and the Human Protein Atlas (CC BY 4.0).^31,32^ **(H)** Cell-cycle programs in each published cell type: share of transcripts and of cycling cells (left), program composition of *E2F1* and *CDC20* transcripts by lineage (middle), and cosine similarity of each program’s gene profile estimated in two lineages (right). **(I)** Program posterior shares of individual tumor epithelial cells in each depth zone. **(J)** Four malignant cells carrying several program classes, with transcripts colored by class; white outline, segmented cell boundary. **(K)** Frequency of program-class combinations among tumor epithelial cells (left) and the number of classes per cell under alternative presence rules (right). **(L)** Microenvironment programs bordering deep tumor, the tumor surface and normal mucosa; bars, combined share of each program group in each region. **(M)** Spearman correlation across 250-µm tiles between tumor program classes and the adjacent microenvironment programs; right, an example pair.

**Figure S4. Tumor-lineage programs are confined to one section, and conserved programs co-occur with them within tumor cells. (A)** Sections as in Fig. 4A; boxes mark the views in B– K, which differ from those of Fig. 4. **(B)** Focused views of the boxed regions in A. **(C)** DAPI image of the views in B. **(D)** Programs specific to the lung tumor, shown in the views of all eight sections; they are confined to the lung section **(E)** Programs specific to the lymph node, as in D **(F)** Programs specific to the ERBB2+ ovarian tumor, as in D. **(G)** Program specific to the prostate tumor, as in D. **(H)** Programs specific to the skin tumor, as in D. **(I)** Programs specific to the breast tumor, as in D. **(J)** Programs specific to the cervical tumor, as in D. **(K)** Program specific to the ERBB2− ovarian tumor, as in D. **(L)** Percentage of segmented cells dominated by a tumor-specific program that also carry the indicated conserved program; numbers below bars, cells carrying both. In the lymph node, B- and T-cell programs take the place of tumor-specific programs. **(M)** One example cell per section (outlined) carrying a tumor-specific and a conserved program from L, with each program’s share of the cell’s transcripts; in the lymph node, a cell carrying the B-cell program.

**Figure S5. Comparisons of inference of multi-sample CRC cancer dataset between Xenium and Visium HD platforms. (A)** Whole-section share of each projected cell type (39 profiles) and program (96) on Xenium (share of molecules) versus the serial Visium HD section (share of UMIs). Solid line, equal share; dashed lines, two-fold difference. Spearman ρ between platforms: cell types 0.87–0.88, programs 0.55–0.65 (P1, P2, P5). **(B)** Log2 ratio of Visium HD to Xenium share (from A) of each projected label against the fraction of its reference profile on the 407 Xenium panel genes used in projection. ρ, Spearman correlation. **(C)** Spatial agreement between the aligned sections by bin size: abundance-weighted mean over labels of the Pearson correlation of each label’s share in shared bins; chance, the same statistic after shifting the Visium HD section. **(D)** Projected scRNA-seq programs (rows) cross-tabulated against the 96 programs learned de novo from the Xenium sections (columns) on the same molecules, pooled over P1, P2 and P5; enrichment, observed over expected under independence. Programs with ≥0.5% of the joint mass are shown. **(E)** As in D, for Visium HD (shares of UMIs). The de novo models of the two platforms are separate fits, so column identifiers in D and E do not correspond.

**Figure S6. Composition shifts explain most expression changes in IPF lung except for a few genes**. **(A)** Change in abundance (share of a core’s molecules) of the published cell types, IPF-more vs control cores: Hodges–Lehmann log₂ shift (dot) with exact 95% confidence interval (line); FDR from two-sided Wilcoxon rank-sum tests. **(B)** As in A, for the FICTURE2 programs. **(C)** Marker genes of the FICTURE2 fibroblast programs (top) and published fibroblast cell types (bottom), control and IPF-more cores pooled. Bars, log₂ ratio of mean share in IPF-more over ctrl cores (not tested); arrowhead, value beyond the axis (+5.1). **(D)** Bulk log₂FC (top) and the percentage of it explained by cell-type (middle) or program (bottom) abundance for the 48 differentially expressed genes of Fig. 6F with |log₂FC| ≥ 1.5 and a program label; segments depend on the order in which labels are added. **(E)** *APLNR* and *TGFB3* per core in segmented cells of each cell type (left) and in molecules of each program (right), for target labels and low-expression comparison labels (grey band); p, two-sided Wilcoxon rank-sum test, IPF-more vs ctrl. Insets, target-to-comparison rate ratio; p, permutation test; program ratios rest on near-zero comparison rates and show direction only.

**Figure S7. Comparison of single-molecule inference by FICTURE2 with FICTURE1 and other methods. (A)** The region of Fig. 7K (FICTURE1 mode) with only the cartilage and maturation programs colored; other programs dark grey. Insets as in Fig. 7G. **(B)** As in A, decoded by FICTURE2. The hypertrophic-chondrocyte program received 94.9% of the region’s Col10a1 transcripts (n = 2,078), against 0.4% in FICTURE1 mode. **(C)** As in A, for the osteoblast and perichondrium programs. **(D)** As in C, decoded by FICTURE2. **(E)** As in A, for the mesenchyme and tendon programs. **(F)** As in E, decoded by FICTURE2. **(G)** As in A, for the vessel, blood and fibroblast programs. **(H)** As in G, decoded by FICTURE2. **(I)** As in A, for the cell-cycle and nuclear-RNA programs. **(J)** As in I, decoded by FICTURE2. The mitosis program received 56.7% of the region’s *Top2a*, *Mki67* and *Cdk1* transcripts (n = 3,912), against 3.4% in FICTURE1 mode. **(K)** Label flows between FICTURE2-XR, FICTURE2-LDA and TopACT for the transcripts of the glomerulus in Fig. 7M–P; percentages are shares of each annotation’s labelled transcripts. **(L)** Labels of endothelial- and mesangial-marker transcripts in five glomeruli. Percentages, share in the matching label (Mix for TopACT, which has no mesangial class); in parentheses, the same share among transcripts that TopACT labelled. **(M)** As in K, for the immune infiltrate of Fig. 7M–P. **(N)** FICTURE2-LDA labels of the marker transcripts of each immune class in the infiltrate; right, share in the matching label. **(O)** Marker-gene enrichment in the labels of FICTURE2-XR, FICTURE2-LDA and TopACT that are shown in Fig. 7M–O, from whole-section pseudobulks. **(P)** Share of immune-marker transcripts in the infiltrate placed in the matching class when each segmented cell or square bin (left, 6 µm) takes its majority marker class, an upper bound for any one-label-per-unit annotation, compared with FICTURE2-LDA; left, per class; right, all markers against unit area.

