## Supplementary Figures 1-7 for "Molecule-resolved inference reveals the spatial organization of transcriptional programs"

### Fig. S1

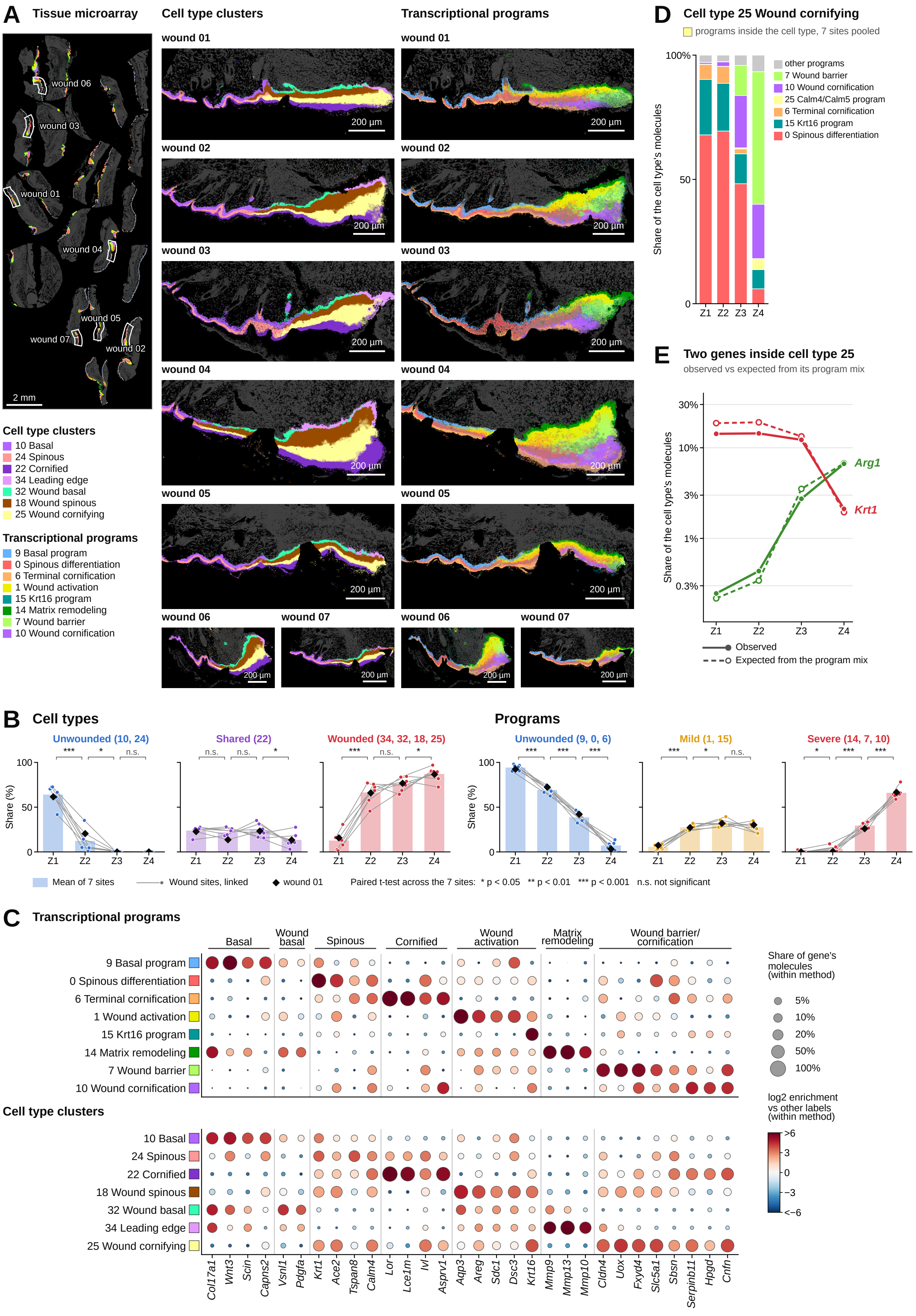

A Tissue microarray
wound 06
wound 03
wound 01
wound 04
wound 05
wound 07
wound 02
2 mm
Cell type clusters
Transcriptional programs
wound 01
wound 02
wound 03
wound 04
wound 05
wound 06
wound 07
200 µm
Cell type clusters
10 Basal
24 Spinous
22 Cornified
34 Leading edge
32 Wound basal
18 Wound spinous
25 Wound cornifying
Transcriptional programs
9 Basal program
0 Spinous differentiation
6 Terminal cornification
1 Wound activation
15 Krt16 program
14 Matrix remodeling
7 Wound barrier
10 Wound cornification
D Cell type 25 Wound cornifying
programs inside the cell type, 7 sites pooled
100%
50
0
Share of the cell type's molecules
Z1 Z2 Z3 Z4
other programs
7 Wound barrier
10 Wound cornification
25 Calm4/Calm5 program
6 Terminal cornification
15 Krt16 program
0 Spinous differentiation
E Two genes inside cell type 25
observed vs expected from its program mix
30%
10%
3%
1%
0.3%
Share of the cell type's molecules
Arg1
Krt1
Z1 Z2 Z3 Z4
Observed
Expected from the program mix
B Cell types
Programs
Unwounded (10, 24)
Shared (22)
Wounded (34, 32, 18, 25)
Unwounded (9, 0, 6)
Mild (1, 15)
Severe (14, 7, 10)
*** * n.s.
n.s. n.s. *
*** n.s. *
*** *** ***
*** * n.s.
* *** ***
100
50
0
Share (%)
Z1 Z2 Z3 Z4
Mean of 7 sites
Wound sites, linked
wound 01
Paired t-test across the 7 sites: * p < 0.05 ** p < 0.01 *** p < 0.001 n.s. not significant
C Transcriptional programs
Basal
Wound basal
Spinous
Cornified
Wound activation
Matrix remodeling
Wound barrier/ cornification
9 Basal program
0 Spinous differentiation
6 Terminal cornification
1 Wound activation
15 Krt16 program
14 Matrix remodeling
7 Wound barrier
10 Wound cornification
Cell type clusters
10 Basal
24 Spinous
22 Cornified
18 Wound spinous
32 Wound basal
34 Leading edge
25 Wound cornifying
Col17a1 Wnt3 Scin Capns2 Vsnl1 Pdgfa Krt1 Ace2 Tspan8 Calm4 Lor Lce1m Ivl Asprv1 Aqp3 Areg Sdc1 Dsc3 Krt16 Mmp9 Mmp13 Mmp10 Cldn4 Uox Fxyd4 Slc5a1 Sbsn Serpinb11 Hpgd Cnfn
Share of gene's molecules (within method)
5%
10%
20%
50%
100%
log2 enrichment vs other labels (within method)
>6
3
0
−3
<−6

### Fig. S2

**A** **Marker genes in the facial muscle**

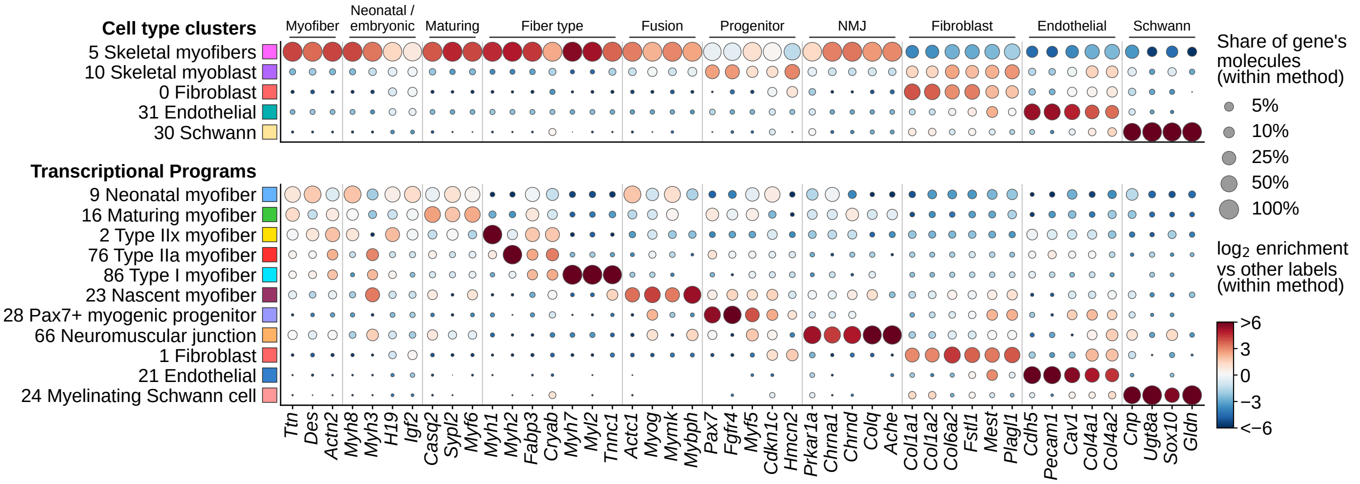


**B** **Molecules of the facial muscle: segmented cells → clusters per pixel → programs**

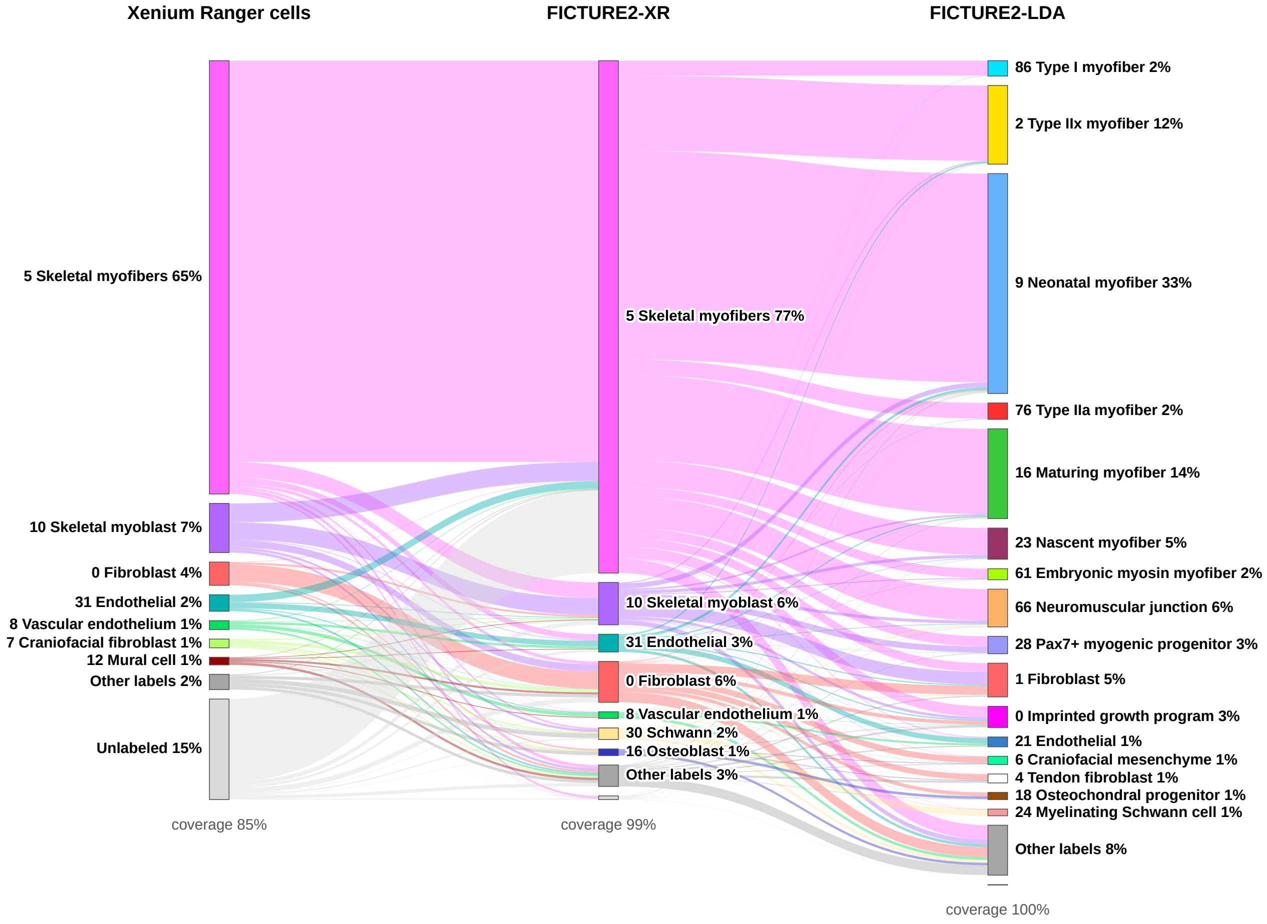

### Fig. S3

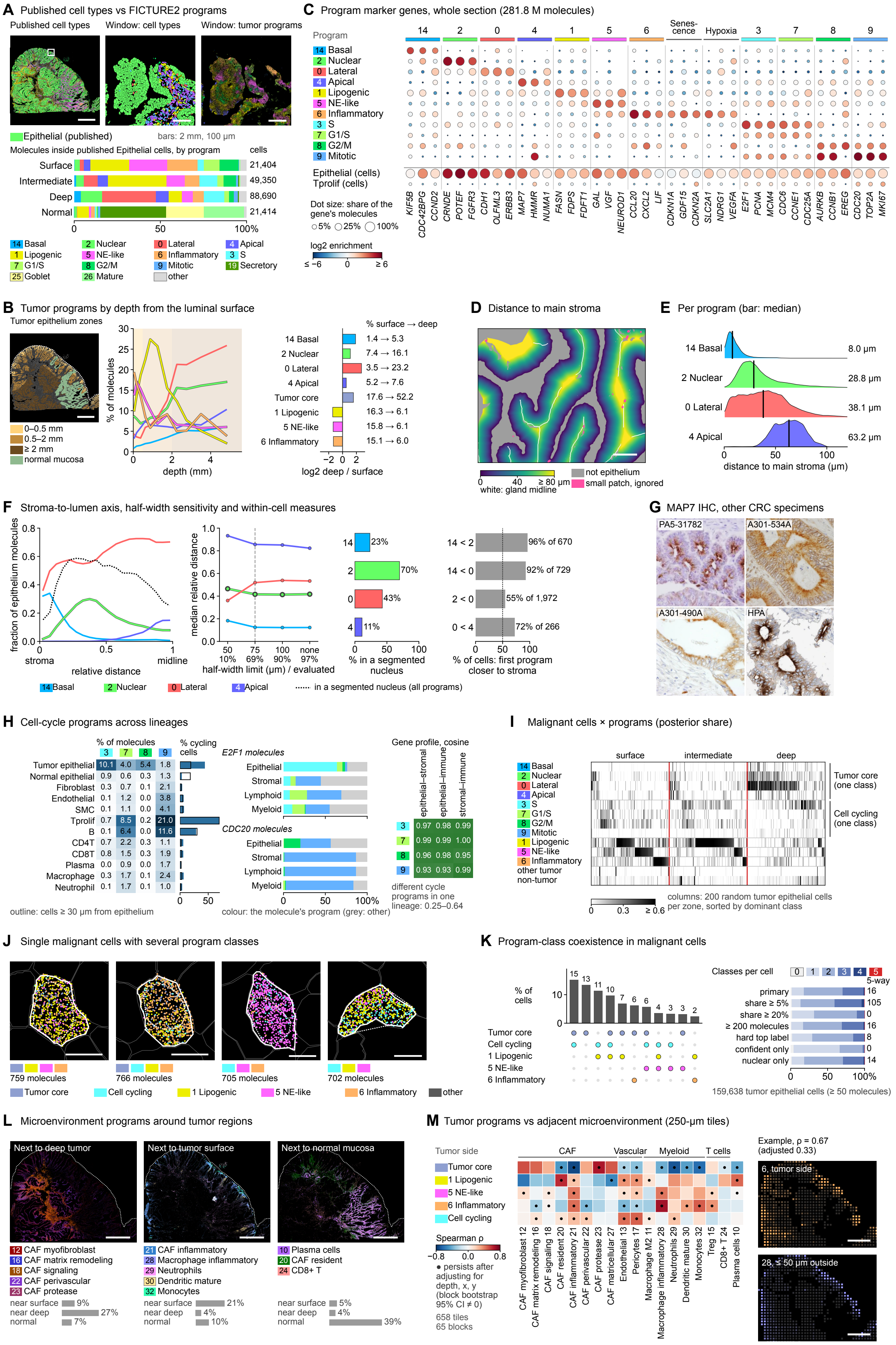
A Published cell types vs FICTURE2 programs
Published cell types
Window: cell types
Window: tumor programs
Epithelial (published)
bars: 2 mm, 100 µm
Molecules inside published Epithelial cells, by program
cells
Surface 21,404
Intermediate 49,350
Deep 88,690
Normal 21,414
0 50 100%
14 Basal
2 Nuclear
0 Lateral
4 Apical
1 Lipogenic
5 NE-like
6 Inflammatory
3 S
7 G1/S
8 G2/M
9 Mitotic
19 Secretory
25 Goblet
26 Mature
other
C Program marker genes, whole section (281.8 M molecules)
Program
14 Basal
2 Nuclear
0 Lateral
4 Apical
1 Lipogenic
5 NE-like
6 Inflammatory
3 S
7 G1/S
8 G2/M
9 Mitotic
Epithelial (cells)
Tprolif (cells)
Dot size: share of the gene's molecules
5% 25% 100%
log2 enrichment
≤ −6 0 ≥ 6
14 2 0 4 1 5 6 Senes-cence Hypoxia 3 7 8 9
KIF5B CDC42BPG CCND2 CRNDE POTEF FGFR3 CDH1 OLFML3 ERBB3 MAP7 HMMR NUMA1 FASN FDPS FDFT1 GAL VGF NEUROD1 CCL20 CXCL2 LIF CDKN1A GDF15 CDKN2A SLC2A1 NDRG1 VEGFA E2F1 PCNA MCM4 CDC6 CCNE1 CDC25A AURKB CCNB1 EREG CDC20 TOP2A MKI67
B Tumor programs by depth from the luminal surface
Tumor epithelium zones
0–0.5 mm
0.5–2 mm
≥ 2 mm
normal mucosa
% of molecules
depth (mm)
% surface → deep
14 Basal 1.4 → 5.3
2 Nuclear 7.4 → 16.1
0 Lateral 3.5 → 23.2
4 Apical 5.2 → 7.6
Tumor core 17.6 → 52.2
1 Lipogenic 16.3 → 6.1
5 NE-like 15.8 → 6.1
6 Inflammatory 15.1 → 6.0
log2 deep / surface
D Distance to main stroma
0 40 ≥ 80 µm
white: gland midline
not epithelium
small patch, ignored
E Per program (bar: median)
14 Basal 8.0 µm
2 Nuclear 28.8 µm
0 Lateral 38.1 µm
4 Apical 63.2 µm
distance to main stroma (µm)
F Stroma-to-lumen axis, half-width sensitivity and within-cell measures
fraction of epithelium molecules
stroma
relative distance
midline
median relative distance
half-width limit (µm) / evaluated
50 75 100 none
10% 69% 90% 97%
14 23%
2 70%
0 43%
4 11%
% in a segmented nucleus
14 < 2 96% of 670
14 < 0 92% of 729
2 < 0 55% of 1,972
0 < 4 72% of 266
% of cells: first program closer to stroma
14 Basal
2 Nuclear
0 Lateral
4 Apical
in a segmented nucleus (all programs)
G MAP7 IHC, other CRC specimens
PA5-31782
A301-534A
A301-490A
HPA
H Cell-cycle programs across lineages
% of molecules
% cycling cells
3 7 8 9
Tumor epithelial 10.1 4.0 5.4 1.8
Normal epithelial 0.9 0.6 0.3 1.3
Fibroblast 0.3 0.7 0.1 2.1
Endothelial 0.1 1.2 0.0 3.8
SMC 0.1 1.1 0.0 4.1
Tprolif 0.7 8.5 0.2 21.0
B 0.1 6.4 0.0 11.6
CD4T 0.7 2.2 0.3 1.1
CD8T 0.8 1.5 0.3 1.9
Plasma 0.0 0.9 0.0 1.7
Macrophage 0.3 1.7 0.1 2.4
Neutrophil 0.1 1.7 0.1 1.0
outline: cells ≥ 30 µm from epithelium
E2F1 molecules
Epithelial
Stromal
Lymphoid
Myeloid
CDC20 molecules
colour: the molecule's program (grey: other)
Gene profile, cosine
epithelial–stromal epithelial–immune stromal–immune
3 0.97 0.98 0.99
7 0.99 0.99 1.00
8 0.96 0.98 0.95
9 0.93 0.93 0.99
different cycle programs in one lineage: 0.25–0.64
I Malignant cells × programs (posterior share)
surface intermediate deep
14 Basal
2 Nuclear
0 Lateral
4 Apical
3 S
7 G1/S
8 G2/M
9 Mitotic
1 Lipogenic
5 NE-like
6 Inflammatory
other tumor
non-tumor
Tumor core (one class)
Cell cycling (one class)
0 0.3 ≥ 0.6
columns: 200 random tumor epithelial cells per zone, sorted by dominant class
J Single malignant cells with several program classes
759 molecules
766 molecules
705 molecules
702 molecules
Tumor core
Cell cycling
1 Lipogenic
5 NE-like
6 Inflammatory
other
K Program-class coexistence in malignant cells
% of cells
15 13 11 10 7 6 6 4 3 3 2
Tumor core
Cell cycling
1 Lipogenic
5 NE-like
6 Inflammatory
Classes per cell
0 1 2 3 4 5-way
primary 16
share ≥ 5% 105
share ≥ 20% 0
≥ 200 molecules 16
hard top label 8
confident only 0
nuclear only 14
159,638 tumor epithelial cells (≥ 50 molecules)
L Microenvironment programs around tumor regions
Next to deep tumor
Next to tumor surface
Next to normal mucosa
12 CAF myofibroblast
16 CAF matrix remodeling
18 CAF signaling
22 CAF perivascular
23 CAF protease
near surface 9%
near deep 27%
normal 7%
21 CAF inflammatory
28 Macrophage inflammatory
29 Neutrophils
30 Dendritic mature
32 Monocytes
near surface 21%
near deep 4%
normal 10%
10 Plasma cells
20 CAF resident
24 CD8+ T
near surface 5%
near deep 4%
normal 39%
M Tumor programs vs adjacent microenvironment (250-µm tiles)
Tumor side
CAF Vascular Myeloid T cells
Tumor core
1 Lipogenic
5 NE-like
6 Inflammatory
Cell cycling
Spearman ρ
−0.8 0.0 0.8
● persists after adjusting for depth, x, y (block bootstrap 95% CI ≠ 0)
658 tiles
65 blocks
CAF myofibroblast 12 CAF matrix remodeling 16 CAF signaling 18 CAF resident 20 CAF inflammatory 21 CAF perivascular 22 CAF protease 23 CAF matricellular 27 Endothelial 13 Pericytes 17 Macrophage M2 11 Macrophage inflammatory 28 Neutrophils 29 Dendritic mature 30 Monocytes 32 Treg 15 CD8+ T 24 Plasma cells 10
Example, ρ = 0.67 (adjusted 0.33)
6, tumor side
28, ≤ 50 µm outside

### Fig. S4

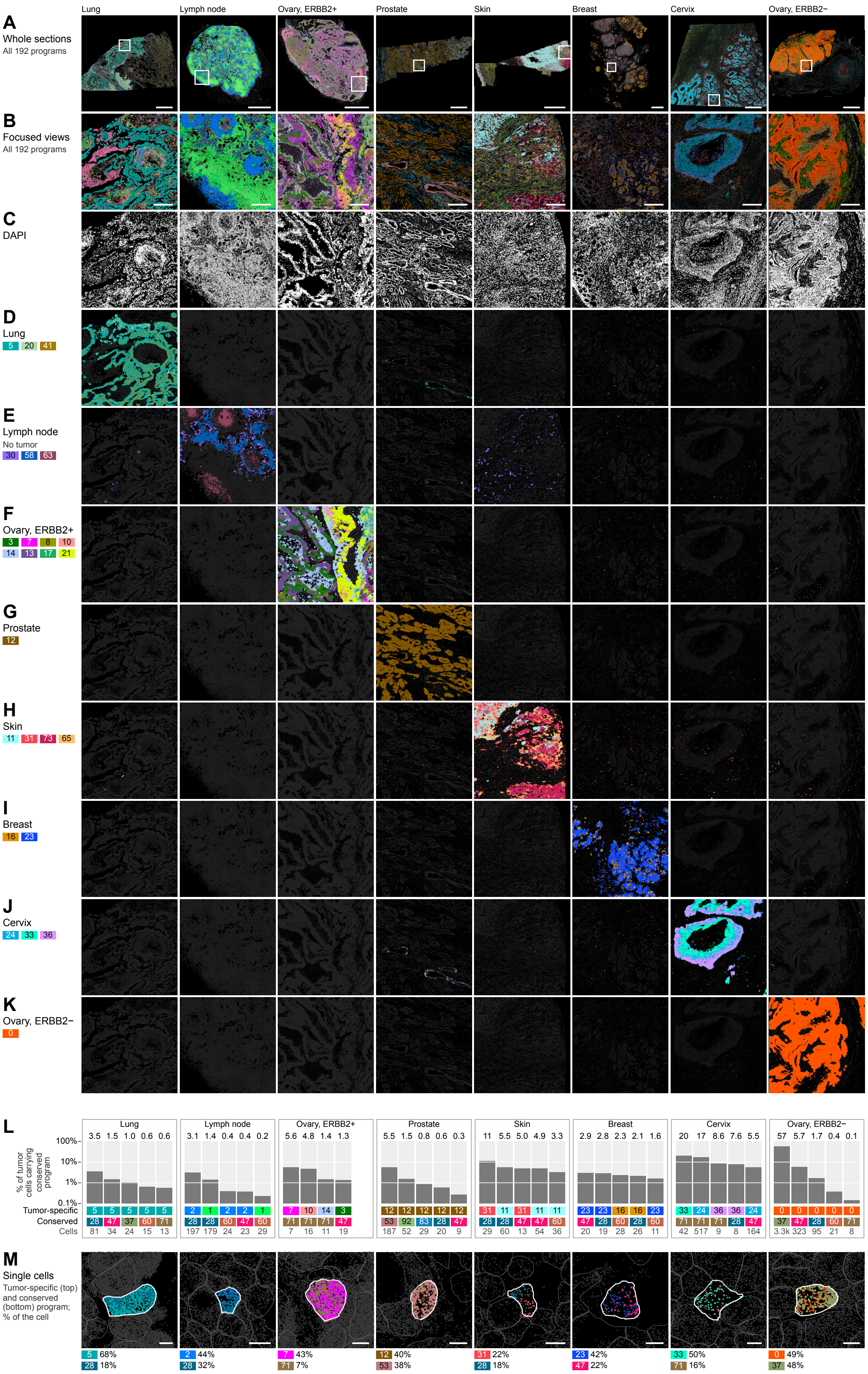
Lung
Lymph node
Ovary, ERBB2+
Prostate
Skin
Breast
Cervix
Ovary, ERBB2−
A
Whole sections
All 192 programs
B
Focused views
All 192 programs
C
DAPI
D
Lung
5 20 41
E
Lymph node
No tumor
30 58 63
F
Ovary, ERBB2+
3 7 8 10
14 13 17 21
G
Prostate
12
H
Skin
11 31 73 65
I
Breast
16 23
J
Cervix
24 33 36
K
Ovary, ERBB2−
0
L
% of tumor cells carrying conserved program
100%
10%
1%
0.1%
Tumor-specific
Conserved
Cells
Lung
3.5 1.5 1.0 0.6 0.6
5 5 5 5 5
28 47 37 60 71
81 34 24 15 13
Lymph node
3.1 1.4 0.4 0.4 0.2
2 1 2 2 1
28 28 60 47 60
197 179 24 23 29
Ovary, ERBB2+
5.6 4.8 1.4 1.3
7 10 14 3
71 71 71 47
7 16 11 19
Prostate
5.5 1.5 0.8 0.6 0.3
12 12 12 12 12
53 92 83 28 47
187 52 29 20 9
Skin
11 5.5 5.0 4.9 3.3
31 11 31 11 11
28 28 47 47 60
29 60 13 54 36
Breast
2.9 2.8 2.3 2.1 1.6
23 23 16 16 23
47 28 60 28 60
20 19 28 26 11
Cervix
20 17 8.6 7.6 5.5
33 24 36 36 24
71 71 71 28 47
42 517 9 8 164
Ovary, ERBB2−
57 5.7 1.7 0.4 0.1
0 0 0 0 0
37 47 28 60 71
3.3k 323 95 21 8
M
Single cells
Tumor-specific (top) and conserved (bottom) program; % of the cell
5 68%
28 18%
2 44%
28 32%
7 43%
71 7%
12 40%
53 38%
31 22%
28 18%
23 42%
47 22%
33 50%
71 16%
0 49%
37 48%

### Fig. S5

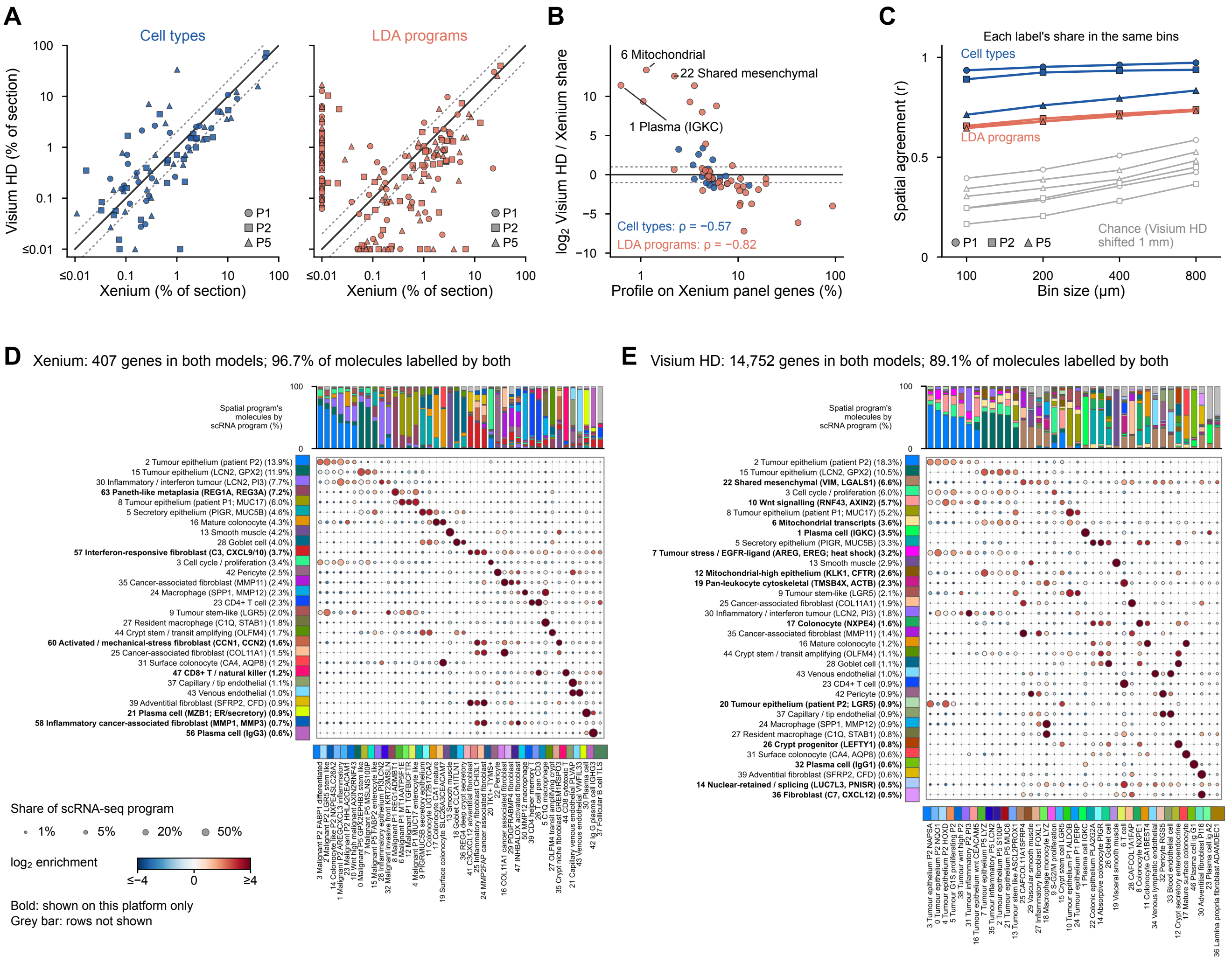
A
Cell types
LDA programs
Visium HD (% of section)
Xenium (% of section)
P1
P2
P5
B
6 Mitochondrial
22 Shared mesenchymal
1 Plasma (IGKC)
log2 Visium HD / Xenium share
Profile on Xenium panel genes (%)
Cell types: ρ = −0.57
LDA programs: ρ = −0.82
C
Each label's share in the same bins
Cell types
LDA programs
Spatial agreement (r)
Chance (Visium HD shifted 1 mm)
Bin size (μm)
D
Xenium: 407 genes in both models; 96.7% of molecules labelled by both
Spatial program's molecules by scRNA program (%)
2 Tumour epithelium (patient P2) (13.9%)
15 Tumour epithelium (LCN2, GPX2) (11.9%)
30 Inflammatory / interferon tumour (LCN2, PI3) (7.7%)
63 Paneth-like metaplasia (REG1A, REG3A) (7.2%)
8 Tumour epithelium (patient P1; MUC17) (6.0%)
5 Secretory epithelium (PIGR, MUC5B) (4.6%)
16 Mature colonocyte (4.3%)
13 Smooth muscle (4.2%)
28 Goblet cell (4.0%)
57 Interferon-responsive fibroblast (C3, CXCL9/10) (3.7%)
3 Cell cycle / proliferation (3.4%)
42 Pericyte (2.5%)
35 Cancer-associated fibroblast (MMP11) (2.4%)
24 Macrophage (SPP1, MMP12) (2.3%)
23 CD4+ T cell (2.3%)
9 Tumour stem-like (LGR5) (2.0%)
27 Resident macrophage (C1Q, STAB1) (1.8%)
44 Crypt stem / transit amplifying (OLFM4) (1.7%)
60 Activated / mechanical-stress fibroblast (CCN1, CCN2) (1.6%)
25 Cancer-associated fibroblast (COL11A1) (1.5%)
31 Surface colonocyte (CA4, AQP8) (1.2%)
47 CD8+ T / natural killer (1.2%)
37 Capillary / tip endothelial (1.1%)
43 Venous endothelial (1.0%)
39 Adventitial fibroblast (SFRP2, CFD) (0.9%)
21 Plasma cell (MZB1; ER/secretory) (0.9%)
58 Inflammatory cancer-associated fibroblast (MMP1, MMP3) (0.7%)
56 Plasma cell (IgG3) (0.6%)
Share of scRNA-seq program
1%
5%
20%
50%
log2 enrichment
≤−4
0
≥4
Bold: shown on this platform only
Grey bar: rows not shown
E
Visium HD: 14,752 genes in both models; 89.1% of molecules labelled by both
Spatial program's molecules by scRNA program (%)
2 Tumour epithelium (patient P2) (18.3%)
15 Tumour epithelium (LCN2, GPX2) (10.5%)
22 Shared mesenchymal (VIM, LGALS1) (6.6%)
3 Cell cycle / proliferation (6.0%)
10 Wnt signalling (RNF43, AXIN2) (5.7%)
8 Tumour epithelium (patient P1; MUC17) (5.2%)
6 Mitochondrial transcripts (3.6%)
1 Plasma cell (IGKC) (3.5%)
5 Secretory epithelium (PIGR, MUC5B) (3.3%)
7 Tumour stress / EGFR-ligand (AREG, EREG; heat shock) (3.2%)
13 Smooth muscle (2.9%)
12 Mitochondrial-high epithelium (KLK1, CFTR) (2.6%)
19 Pan-leukocyte cytoskeletal (TMSB4X, ACTB) (2.3%)
9 Tumour stem-like (LGR5) (2.1%)
25 Cancer-associated fibroblast (COL11A1) (1.9%)
30 Inflammatory / interferon tumour (LCN2, PI3) (1.8%)
17 Colonocyte (NXPE4) (1.6%)
35 Cancer-associated fibroblast (MMP11) (1.4%)
16 Mature colonocyte (1.2%)
44 Crypt stem / transit amplifying (OLFM4) (1.1%)
28 Goblet cell (1.1%)
43 Venous endothelial (1.0%)
23 CD4+ T cell (0.9%)
42 Pericyte (0.9%)
20 Tumour epithelium (patient P2; LGR5) (0.9%)
37 Capillary / tip endothelial (0.9%)
24 Macrophage (SPP1, MMP12) (0.9%)
27 Resident macrophage (C1Q, STAB1) (0.8%)
26 Crypt progenitor (LEFTY1) (0.8%)
31 Surface colonocyte (CA4, AQP8) (0.6%)
32 Plasma cell (IgG1) (0.6%)
39 Adventitial fibroblast (SFRP2, CFD) (0.6%)
14 Nuclear-retained / splicing (LUC7L3, PNISR) (0.5%)
36 Fibroblast (C7, CXCL12) (0.5%)

### Fig. S6

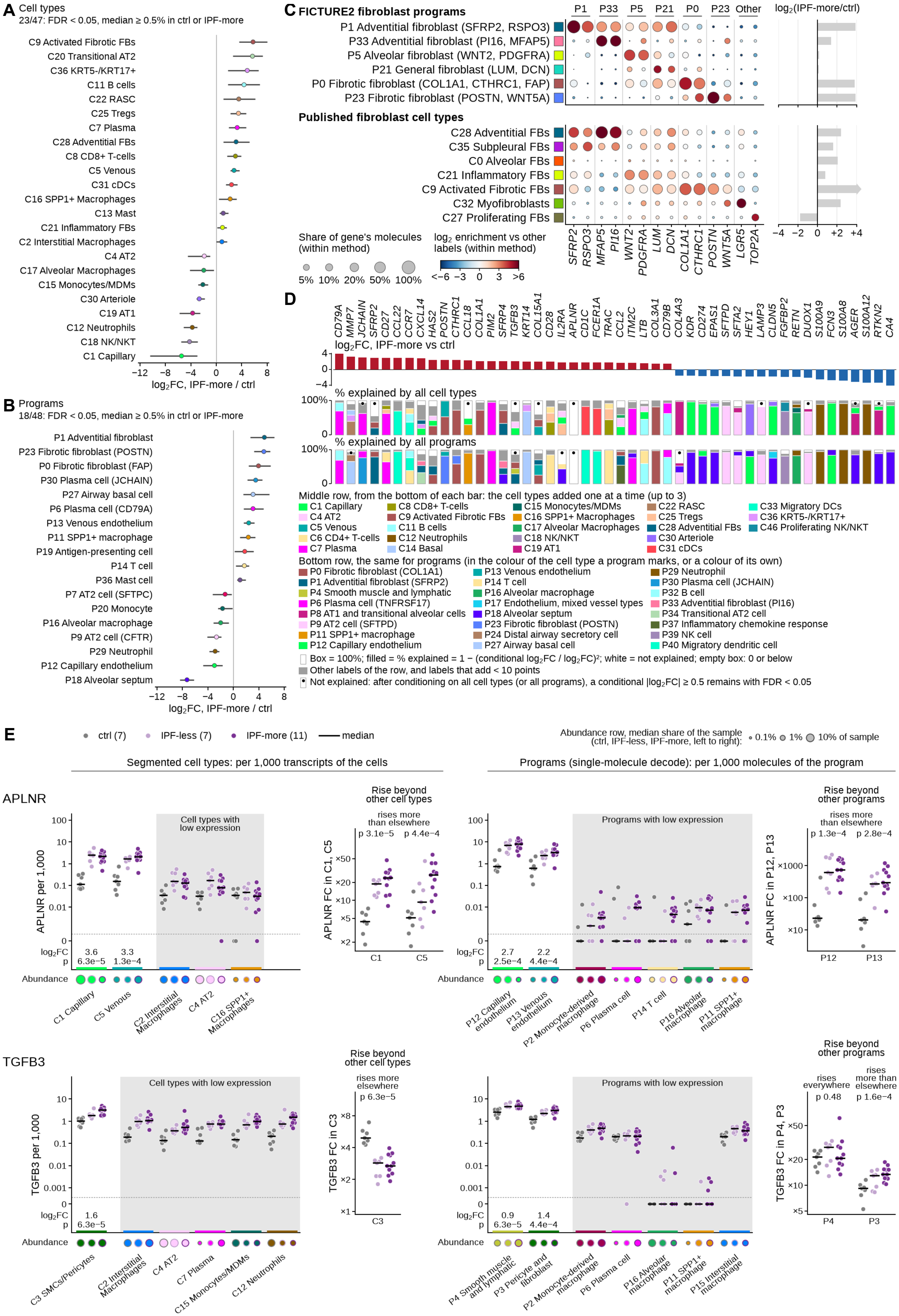

A Cell types
23/47: FDR < 0.05, median ≥ 0.5% in ctrl or IPF-more
C9 Activated Fibrotic FBs
C20 Transitional AT2
C36 KRT5-/KRT17+
C11 B cells
C22 RASC
C25 Tregs
C7 Plasma
C28 Adventitial FBs
C8 CD8+ T-cells
C5 Venous
C31 cDCs
C16 SPP1+ Macrophages
C13 Mast
C21 Inflammatory FBs
C2 Interstitial Macrophages
C4 AT2
C17 Alveolar Macrophages
C15 Monocytes/MDMs
C30 Arteriole
C19 AT1
C12 Neutrophils
C18 NK/NKT
C1 Capillary
−12 −8 −4 0 4 8
log2FC, IPF-more / ctrl
B Programs
18/48: FDR < 0.05, median ≥ 0.5% in ctrl or IPF-more
P1 Adventitial fibroblast
P23 Fibrotic fibroblast (POSTN)
P0 Fibrotic fibroblast (FAP)
P30 Plasma cell (JCHAIN)
P27 Airway basal cell
P6 Plasma cell (CD79A)
P13 Venous endothelium
P11 SPP1+ macrophage
P19 Antigen-presenting cell
P14 T cell
P36 Mast cell
P7 AT2 cell (SFTPC)
P20 Monocyte
P16 Alveolar macrophage
P9 AT2 cell (CFTR)
P29 Neutrophil
P12 Capillary endothelium
P18 Alveolar septum
−12 −8 −4 0 4 8
log2FC, IPF-more / ctrl
C FICTURE2 fibroblast programs
P1 P33 P5 P21 P0 P23 Other
log2(IPF-more/ctrl)
P1 Adventitial fibroblast (SFRP2, RSPO3)
P33 Adventitial fibroblast (PI16, MFAP5)
P5 Alveolar fibroblast (WNT2, PDGFRA)
P21 General fibroblast (LUM, DCN)
P0 Fibrotic fibroblast (COL1A1, CTHRC1, FAP)
P23 Fibrotic fibroblast (POSTN, WNT5A)
Published fibroblast cell types
C28 Adventitial FBs
C35 Subpleural FBs
C0 Alveolar FBs
C21 Inflammatory FBs
C9 Activated Fibrotic FBs
C32 Myofibroblasts
C27 Proliferating FBs
Share of gene's molecules (within method)
5% 10% 20% 50% 100%
log2 enrichment vs other labels (within method)
<−6 −3 0 3 >6
SFRP2 RSPO3 MFAP5 PI16 WNT2 PDGFRA LUM DCN COL1A1 CTHRC1 POSTN WNT5A LGR5 TOP2A
−4 −2 0 +2 +4
D
CD79A MMP7 JCHAIN SFRP2 CD27 CCL22 CCR7 CXCL14 HAS2 POSTN CTHRC1 CCL18 COL1A1 PIM2 SFRP4 TGFB3 KRT14 COL15A1 CD28 IL2RA APLNR CD1C FCER1A TRAC CCL2 ITM2C LTB COL3A1 CD79B COL4A3 KDR CD274 EPAS1 SFTPD SFTA2 HEY1 LAMP3 CLDN5 FGFBP2 RETN DUOX1 S100A9 FCN3 S100A8 AGER S100A12 RTKN2 CA4
log2FC, IPF-more vs ctrl
4 0 −4
% explained by all cell types
100% 0
% explained by all programs
100% 0
Middle row, from the bottom of each bar: the cell types added one at a time (up to 3)
C1 Capillary
C4 AT2
C5 Venous
C6 CD4+ T-cells
C7 Plasma
C8 CD8+ T-cells
C9 Activated Fibrotic FBs
C11 B cells
C12 Neutrophils
C14 Basal
C15 Monocytes/MDMs
C16 SPP1+ Macrophages
C17 Alveolar Macrophages
C18 NK/NKT
C19 AT1
C22 RASC
C25 Tregs
C28 Adventitial FBs
C30 Arteriole
C31 cDCs
C33 Migratory DCs
C36 KRT5-/KRT17+
C46 Proliferating NK/NKT
Bottom row, the same for programs (in the colour of the cell type a program marks, or a colour of its own)
P0 Fibrotic fibroblast (COL1A1)
P1 Adventitial fibroblast (SFRP2)
P4 Smooth muscle and lymphatic
P6 Plasma cell (TNFRSF17)
P8 AT1 and transitional alveolar cells
P9 AT2 cell (SFTPD)
P11 SPP1+ macrophage
P12 Capillary endothelium
P13 Venous endothelium
P14 T cell
P16 Alveolar macrophage
P17 Endothelium, mixed vessel types
P18 Alveolar septum
P23 Fibrotic fibroblast (POSTN)
P24 Distal airway secretory cell
P27 Airway basal cell
P29 Neutrophil
P30 Plasma cell (JCHAIN)
P32 B cell
P33 Adventitial fibroblast (PI16)
P34 Transitional AT2 cell
P37 Inflammatory chemokine response
P39 NK cell
P40 Migratory dendritic cell
Box = 100%; filled = % explained = 1 − (conditional log2FC / log2FC)²; white = not explained; empty box: 0 or below
Other labels of the row, and labels that add < 10 points
Not explained: after conditioning on all cell types (or all programs), a conditional |log2FC| ≥ 0.5 remains with FDR < 0.05
E
ctrl (7) IPF-less (7) IPF-more (11) median
Abundance row, median share of the sample (ctrl, IPF-less, IPF-more, left to right): 0.1% 1% 10% of sample
Segmented cell types: per 1,000 transcripts of the cells
Programs (single-molecule decode): per 1,000 molecules of the program
APLNR
APLNR per 1,000
100 10 1 0.1 0.01 0
Cell types with low expression
log2FC 3.6 3.3
p 6.3e−5 1.3e−4
Abundance
C1 Capillary
C5 Venous
C2 Interstitial Macrophages
C4 AT2
C16 SPP1+ Macrophages
Rise beyond other cell types
rises more than elsewhere
p 3.1e−5 p 4.4e−4
APLNR FC in C1, C5
×50 ×20 ×10 ×5 ×2
C1 C5
Programs with low expression
log2FC 2.7 2.2
p 2.5e−4 4.4e−4
P12 Capillary endothelium
P13 Venous endothelium
P2 Monocyte-derived macrophage
P6 Plasma cell
P14 T cell
P16 Alveolar macrophage
P11 SPP1+ macrophage
Rise beyond other programs
rises more than elsewhere
p 1.3e−4 p 2.8e−4
APLNR FC in P12, P13
×1000 ×100 ×10
P12 P13
TGFB3
TGFB3 per 1,000
10 1 0.1 0.01 0.001 0
Cell types with low expression
log2FC 1.6
p 6.3e−5
Abundance
C3 SMCs/Pericytes
C2 Interstitial Macrophages
C4 AT2
C7 Plasma
C15 Monocytes/MDMs
C12 Neutrophils
Rise beyond other cell types
rises more elsewhere
p 6.3e−5
TGFB3 FC in C3
×8 ×4 ×2 ×1
C3
Programs with low expression
log2FC 0.9 1.4
p 6.3e−5 4.4e−4
P4 Smooth muscle and lymphatic
P3 Pericyte and fibroblast
P2 Monocyte-derived macrophage
P6 Plasma cell
P16 Alveolar macrophage
P11 SPP1+ macrophage
P15 Interstitial macrophage
Rise beyond other programs
rises everywhere
p 0.48
rises more than elsewhere
p 1.6e−4
TGFB3 FC in P4, P3
×50 ×20 ×10 ×5
P4 P3

### Fig. S7

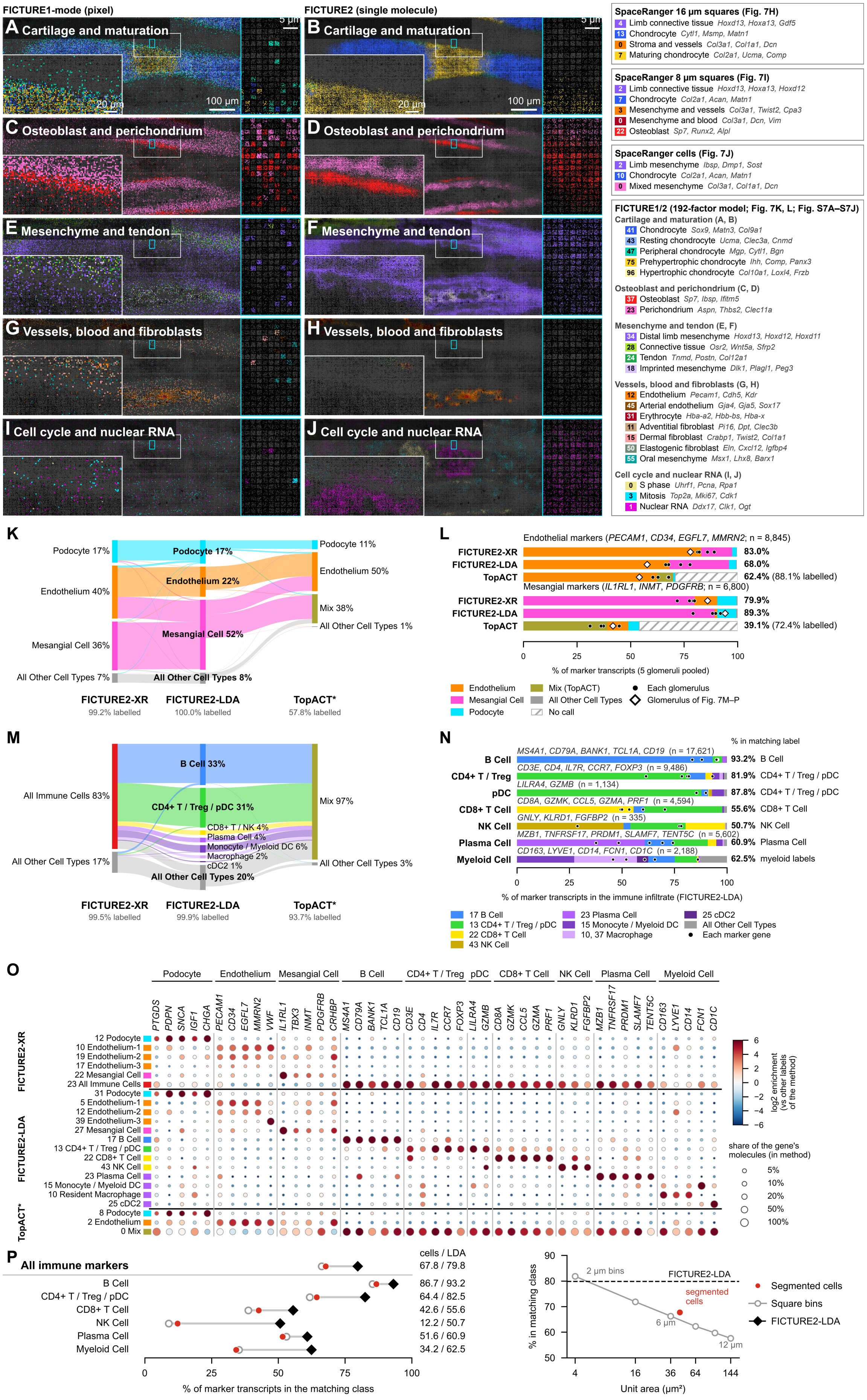

FIGURE1-mode (pixel)
FIGURE2 (single molecule)
A Cartilage and maturation
B Cartilage and maturation
C Osteoblast and perichondrium
D Osteoblast and perichondrium
E Mesenchyme and tendon
F Mesenchyme and tendon
G Vessels, blood and fibroblasts
H Vessels, blood and fibroblasts
I Cell cycle and nuclear RNA
J Cell cycle and nuclear RNA
5 µm
20 µm
100 µm
SpaceRanger 16 µm squares (Fig. 7H)
4 Limb connective tissue Hoxd13, Hoxa13, Gdf5
13 Chondrocyte Cytl1, Msmp, Matn1
0 Stroma and vessels Col3a1, Col1a1, Dcn
7 Maturing chondrocyte Col2a1, Ucma, Comp
SpaceRanger 8 µm squares (Fig. 7I)
2 Limb connective tissue Hoxd13, Hoxa13, Hoxd12
7 Chondrocyte Col2a1, Acan, Matn1
3 Mesenchyme and vessels Col3a1, Twist2, Cpa3
0 Mesenchyme and blood Col3a1, Dcn, Vim
22 Osteoblast Sp7, Runx2, Alpl
SpaceRanger cells (Fig. 7J)
2 Limb mesenchyme Ibsp, Dmp1, Sost
10 Chondrocyte Col2a1, Acan, Matn1
0 Mixed mesenchyme Col3a1, Col1a1, Dcn
FICTURE1/2 (192-factor model; Fig. 7K, L; Fig. S7A–S7J)
Cartilage and maturation (A, B)
41 Chondrocyte Sox9, Matn3, Col9a1
43 Resting chondrocyte Ucma, Clec3a, Cnmd
47 Peripheral chondrocyte Mgp, Cytl1, Bgn
75 Prehypertrophic chondrocyte Ihh, Comp, Panx3
96 Hypertrophic chondrocyte Col10a1, Loxl4, Frzb
Osteoblast and perichondrium (C, D)
37 Osteoblast Sp7, Ibsp, Ifitm5
23 Perichondrium Aspn, Thbs2, Clec11a
Mesenchyme and tendon (E, F)
34 Distal limb mesenchyme Hoxd13, Hoxd12, Hoxd11
28 Connective tissue Osr2, Wnt5a, Sfrp2
24 Tendon Tnmd, Postn, Col12a1
18 Imprinted mesenchyme Dlk1, Plagl1, Peg3
Vessels, blood and fibroblasts (G, H)
12 Endothelium Pecam1, Cdh5, Kdr
45 Arterial endothelium Gja4, Gja5, Sox17
31 Erythrocyte Hba-a2, Hbb-bs, Hba-x
11 Adventitial fibroblast Pi16, Dpt, Clec3b
15 Dermal fibroblast Crabp1, Twist2, Col1a1
50 Elastogenic fibroblast Eln, Cxcl12, Igfbp4
55 Oral mesenchyme Msx1, Lhx8, Barx1
Cell cycle and nuclear RNA (I, J)
0 S phase Uhrf1, Pcna, Rpa1
3 Mitosis Top2a, Mki67, Cdk1
1 Nuclear RNA Ddx17, Clk1, Ogt
K
Podocyte 17%
Endothelium 40%
Mesangial Cell 36%
All Other Cell Types 7%
Podocyte 17%
Endothelium 22%
Mesangial Cell 52%
All Other Cell Types 8%
Podocyte 11%
Endothelium 50%
Mix 38%
All Other Cell Types 1%
FICTURE2-XR
99.2% labelled
FICTURE2-LDA
100.0% labelled
TopACT*
57.8% labelled
L
Endothelial markers (PECAM1, CD34, EGFL7, MMRN2; n = 8,845)
FICTURE2-XR 83.0%
FICTURE2-LDA 68.0%
TopACT 62.4% (88.1% labelled)
Mesangial markers (IL1RL1, INMT, PDGFRB; n = 6,800)
FICTURE2-XR 79.9%
FICTURE2-LDA 89.3%
TopACT 39.1% (72.4% labelled)
0 25 50 75 100
% of marker transcripts (5 glomeruli pooled)
Endothelium
Mesangial Cell
Podocyte
Mix (TopACT)
All Other Cell Types
No call
Each glomerulus
Glomerulus of Fig. 7M–P
M
All Immune Cells 83%
All Other Cell Types 17%
B Cell 33%
CD4+ T / Treg / pDC 31%
CD8+ T / NK 4%
Plasma Cell 4%
Monocyte / Myeloid DC 6%
Macrophage 2%
cDC2 1%
All Other Cell Types 20%
Mix 97%
All Other Cell Types 3%
FICTURE2-XR
99.5% labelled
FICTURE2-LDA
99.9% labelled
TopACT*
93.7% labelled
N
% in matching label
B Cell MS4A1, CD79A, BANK1, TCL1A, CD19 (n = 17,621) 93.2% B Cell
CD4+ T / Treg CD3E, CD4, IL7R, CCR7, FOXP3 (n = 9,486) 81.9% CD4+ T / Treg / pDC
pDC LILRA4, GZMB (n = 1,134) 87.8% CD4+ T / Treg / pDC
CD8+ T Cell CD8A, GZMK, CCL5, GZMA, PRF1 (n = 4,594) 55.6% CD8+ T Cell
NK Cell GNLY, KLRD1, FGFBP2 (n = 335) 50.7% NK Cell
Plasma Cell MZB1, TNFRSF17, PRDM1, SLAMF7, TENT5C (n = 5,602) 60.9% Plasma Cell
Myeloid Cell CD163, LYVE1, CD14, FCN1, CD1C (n = 2,188) 62.5% myeloid labels
0 25 50 75 100
% of marker transcripts in the immune infiltrate (FICTURE2-LDA)
17 B Cell
13 CD4+ T / Treg / pDC
22 CD8+ T Cell
43 NK Cell
23 Plasma Cell
15 Monocyte / Myeloid DC
10, 37 Macrophage
25 cDC2
All Other Cell Types
Each marker gene
O
Podocyte
Endothelium
Mesangial Cell
B Cell
CD4+ T / Treg
pDC
CD8+ T Cell
NK Cell
Plasma Cell
Myeloid Cell
PTGDS PDPN SNCA IGF1 CHGA PECAM1 CD34 EGFL7 MMRN2 VWF IL1RL1 TBX3 INMT PDGFRB CRHBP MS4A1 CD79A BANK1 TCL1A CD19 CD3E CD4 IL7R CCR7 FOXP3 LILRA4 GZMB CD8A GZMK CCL5 GZMA PRF1 GNLY KLRD1 FGFBP2 MZB1 TNFRSF17 PRDM1 SLAMF7 TENT5C CD163 LYVE1 CD14 FCN1 CD1C
FICTURE2-XR
12 Podocyte
10 Endothelium-1
19 Endothelium-2
17 Endothelium-3
22 Mesangial Cell
23 All Immune Cells
FICTURE2-LDA
31 Podocyte
5 Endothelium-1
12 Endothelium-2
39 Endothelium-3
27 Mesangial Cell
17 B Cell
13 CD4+ T / Treg / pDC
22 CD8+ T Cell
43 NK Cell
23 Plasma Cell
15 Monocyte / Myeloid DC
10 Resident Macrophage
25 cDC2
TopACT*
8 Podocyte
2 Endothelium
0 Mix
log2 enrichment (vs other labels of the method)
6 4 2 0 −2 −4 −6
share of the gene's molecules (in method)
5%
10%
20%
50%
100%
P
cells / LDA
All immune markers 67.8 / 79.8
B Cell 86.7 / 93.2
CD4+ T / Treg / pDC 64.4 / 82.5
CD8+ T Cell 42.6 / 55.6
NK Cell 12.2 / 50.7
Plasma Cell 51.6 / 60.9
Myeloid Cell 34.2 / 62.5
0 25 50 75 100
% of marker transcripts in the matching class
90 80 70 60 50
% in matching class
2 µm bins
FICTURE2-LDA
segmented cells
6 µm
12 µm
4 16 36 64 144
Unit area (µm²)
Segmented cells
Square bins
FICTURE2-LDA
